# Low-density lipoprotein receptor-targeting chimeras for membrane protein degradation and enhanced drug delivery

**DOI:** 10.1101/2025.06.06.658366

**Authors:** Fangzhu Zhao, Yan Wu, Kaitlin Schaefer, Yun Zhang, Kun Miao, Zi Yao, Snehal D. Ganjave, Kaan Kumru, Trenton M. Peters-Clarke, Alex Inague, James A. Olzmann, Kevin K. Leung, James A. Wells

## Abstract

Antibody-based therapeutics encompass diverse modalities for targeting tumor cells. Among these, antibody-drug conjugates (ADCs) and extracellular targeted protein degradation (eTPD) specifically depend on efficient lysosomal trafficking for activity. However, many tumor antigens exhibit poor internalization, limiting ADC effectiveness. To address this, we developed low-density lipoprotein receptor-targeting chimeras (LIPTACs), leveraging the constitutive endocytic and recycling activity of the LDLR to enhance lysosomal delivery. LIPTACs enable efficient and selective degradation of diverse extracellular membrane proteins. Additionally, by coupling LIPTACs with cytotoxic payloads to generate degrader-drug conjugates, we can achieve superior intracellular delivery and enhanced cytotoxicity compared to conventional ADCs. The dual modality addresses key challenges of inadequate internalization in conventional ADCs and cytotoxic potency for current eTPD strategies. Our findings demonstrate that LDLR-mediated trafficking can enhance eTPD and ADCs, providing a hybrid blueprint for developing next-generation antibody therapeutics with broader utility and improved efficacy in cancer treatment.

## Introduction

Extracellular and membrane-associated proteins represent approximately one-third of all protein-coding genes and are key targets for antibody-based therapeutics^1^. Antibodies provide diverse mechanisms of action in cancer therapy, including receptor inhibition or activation^2^, immune cell recruitment^3^, antibody drug conjugates (ADCs) for targeted toxin delivery^4^, and extracellular targeted protein degradation (eTPD) for proteolytic degradation^5^. Among these, ADCs combine tumor-targeting antibodies with cytotoxic drugs to achieve selective tumor cell killing^6^. However, the efficacy of ADCs hinges on the efficient internalization of the antibody-antigen complex to facilitate intracellular drug release in the lysosome. Not all surface antigens undergo productive internalization upon antibody binding, posing a significant limitation to ADC effectiveness^7^. Recent innovations in bispecific antibodies have sought to overcome this hurdle by enhancing receptor internalization through receptor clustering, such as biparatopic ADCs^8,9^, or by targeting fast-internalizing receptors in dual-antigen strategies for more efficient lysosomal trafficking^10,11^.

In parallel, eTPD has emerged as a promising therapeutic approach that co-opts natural endolysosomal pathways to selectively degrade membrane-bound and soluble extracellular proteins. Unlike intracellular targeted protein degradation (iTPD) that is driven by recruiting the proteasome for degradation, eTPD mostly shuttles extracellular proteins to the lysosome for degradation. Furthermore, whereas iTPD generally uses only two different widely expressed E3 ligases^12^ cereblon and von Hippel–Lindau (VHL), eTPD utilizes a wide array of cell surface degrader systems including: neonatal Fc receptor (FcRn)^13^, glycan binding receptors^14,15^, transmembrane E3 ligases^16–18^, cytokine receptors^19^, integrins^20^, and transferrin receptors^21^. Increasing the optionality of cell surface degraders offers greater opportunity for cell specific eTPD. Moreover, ADCs with cleavable linkers require the same lysosomal trafficking system as eTPD. To this end, we wondered if it was possible to develop degrader–drug conjugates (DDCs)—a new class of bifunctional therapeutics that intentionally hybridizes eTPD with ADC for greater efficiency of drug payload delivery.

To implement this approach, we targeted the robust recycling low-density lipoprotein receptor (LDLR) as a lysosomal trafficking effector. LDLR naturally internalizes LDL via clathrin-mediated endocytosis^22,23^ and facilitates its delivery to the lysosome at low pH^24^. LDLR is upregulated in proliferating cancer cells^25,26^ and activated T cells^27^. It is one of the fastest and most efficient internalizers that recycles through the lysosome every 12 minutes^28^. These features make the LDLR an attractive candidate for eTPD we refer to as LDLR-targeting chimeras (LIPTACs). We show that LIPTACs mediate selective and efficient lysosomal degradation for multiple membrane proteins. Furthermore, by conjugating cytotoxic payloads to LIPTACs or cytokine receptor-targeting chimeras (KineTACs)^19^, we show these DDCs can boost the potencies of conventional ADCs by up to 20-fold. This work highlights LIPTACs as a versatile platform to enhance payload delivery and broaden the therapeutic utility of antibody-based modalities.

## Results

### Selection and characterization of LDLR antibodies

The extracellular portion of the LDLR contains the ligand binding domain, the epidermal growth factor (EGF)-like domain, and the O-link sugar domain^29^. We and others have observed that the ligand-binding domain of the LDLR can be shed to varying degrees in cells^30,31^ transformed with *KRAS(G12V)* or *HER2*. This prompted us to select for antibodies against the membrane proximal EGF-like domain of the LDLR in order to preserve its ligand ability for LDL uptake and to enable recruitment of both full-length and cleaved forms (cLDLR) for eTPD. After four rounds of phage selection, we plated 96 single phage colonies for screening using an enzyme-linked immunosorbent assay (ELISA) (**Extended Data Fig.1a**,**1b**).

Phage that passed initial screening were expressed recombinantly as monoclonal fragment antigen-binding (Fab) antibodies for further characterization. The ELISA binding assay against the cLDLR or full-length (flLDLR) identified multiple high affinity binding-affinity clones (**Fig. 1a**, **Extended Data Fig.2a**) such as 142F1 and 142F6. Flow cytometry confirmed that all Fabs bound to LDLR^+^ MDA-MB-231 cells (**Fig. 1b**). None of the Fabs exhibited cross-reactivity with the other members of the LDLR family, including the LDLR-related protein 2 (LRP2), LRP8 and VLDLR (**Extended Data Fig.2a**). Additionally, the Fabs displayed minimal polyreactive binding across the nonspecific antigen panels^32^ (**Extended Data Fig.2b**). Epitope binning experiments by biolayer interferometry (BLI) showed Fabs 142F1 and 142F6 could bind simultaneously, indicating two distinct and non-competitive epitopes (**Fig. 1c**, **Extended Data Fig.3**). BLI experiments showed that 142F1 and 142F6 bound to cLDLR with affinities of 5.8 nM and 16 nM, respectively (**Fig. 1d**).

**Fig. 1.**
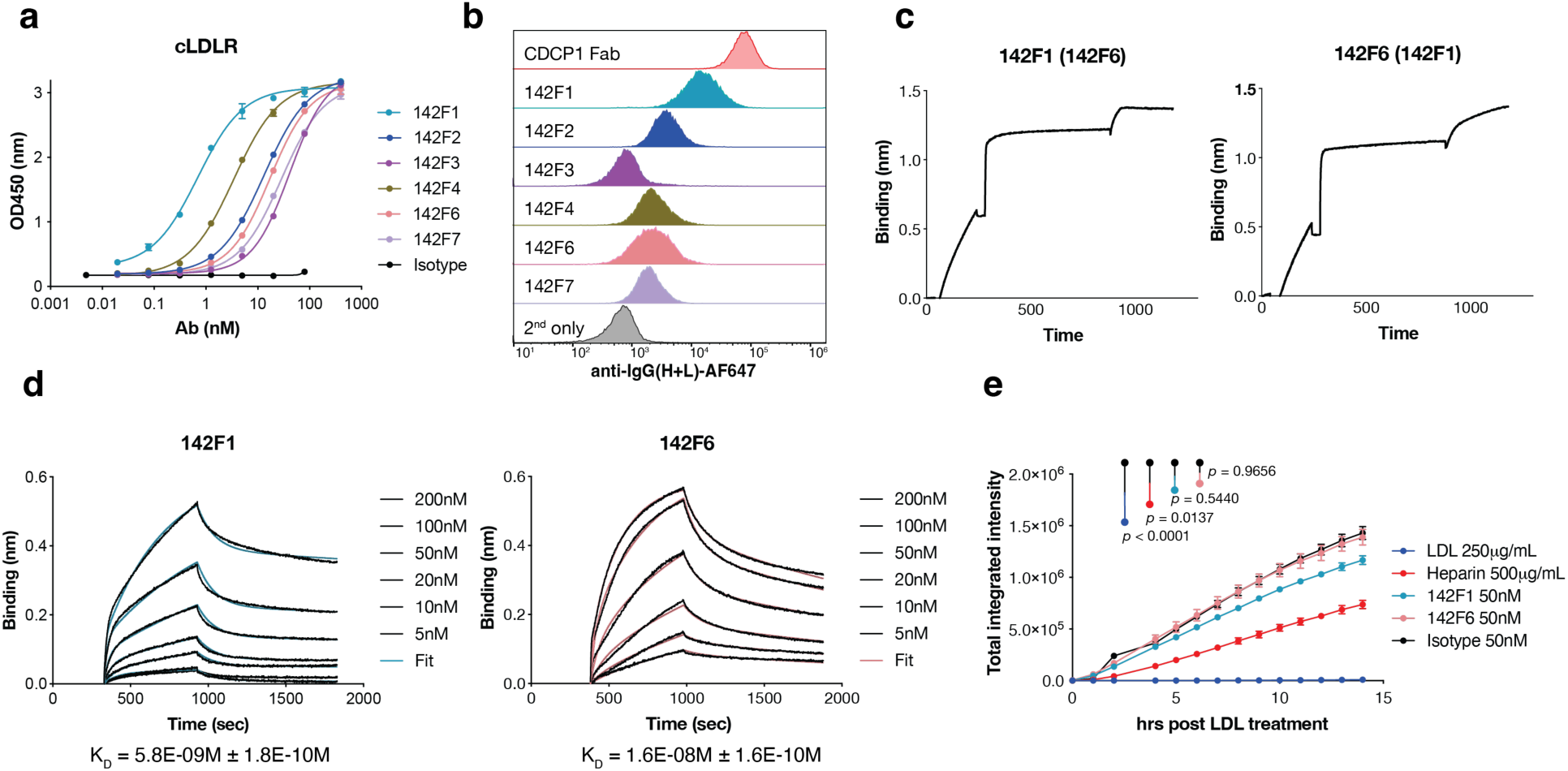
Characterization of cLDLR-specific antibodies. **a**, ELISA binding of recombinant Fabs against the cLDLR antigen. Absorbance was read at 450 nm. An anti-CDCP1 Fab 4A06^54^ was used as a negative isotype control. Each sample was tested in biological duplicate and error bars represented standard deviations. **b**, Flow cytometry of different Fabs binding to LDLR^+^ MDA-MB-231 cells. 50 nM of each Fab was incubated with cells for 30 min and washed twice, followed by AF647 conjugated goat anti-human IgG (H+L) antibody staining for 15 min. **c**, Epitope binning of two anti-LDLR Fabs, 142F1 and 142F6 revealed two different epitopes on cLDLR. Biotinylated cLDLR was captured using a streptavidin biosensor and indicated antibodies at a concentration of 200 nM were incubated for 10 min followed by incubation with 50 nM of the second competing antibodies for 5 min. **d**, BLI analysis of 142F1 and 142F6 Fabs to estimate their affinities to cLDLR. Biontinylated cLDLR was immobilized via the streptavidin biosensor and varying concentrations of each Fab was injected. Black lines were the experimental trace obtained from the BLI experiments and colored lines were the global fits. **e**, Internalization of pH-sensitive Phrodo red-labeled LDL on HeLa cells after 30 min pretreatment with LDL, heparin, or each Fab, respectively. Total integrated intensity is calculated by ROCU x μm^2^/image on the Incucyte software. Each sample was tested in biological triplicate and error bars represent standard deviations. Statistics were calculated by one-way ANOVA and Holm-Sidak multiple comparisons test.

Given that LDLR is essential for regulating plasma cholesterol levels, we then investigated whether our LDLR Fabs affected LDL uptake. We used an Incuyte-based uptake assay with fluorescently labeled (pHrodo) LDL to monitor trafficking to low pH vesicles. We serum starved Hela cells, treated them with LDLR Fabs, heparin, or unlabeled competitor LDL, and then added pHrodo red dye-labeled LDL. As expected, pHrodo red dye-labeled LDL trafficked robustly in both the presence and absence of LDLR Fabs, but was inhibited upon addition of unlabeled LDL and heparin^33^ (**Fig. 1e**). These data suggest the LDLR-mediated LDL trafficking remains intact upon Fab binding.

### Design of LIPTAC degraders for EGFR degradation

The design of LIPTACs involved a bispecific antibody, with one arm targeting a protein of interest (POI) and the other arm recruiting LDLR to bring the POI and LDLR in close proximity for lysosomal trafficking and eTPD. As a proof-of-concept, we generated LIPTACs to degrade EGF receptor (EGFR), a receptor tyrosine kinase that plays a critical role in the development and progression of various types of cancers^34,35^. The therapeutic anti-EGFR Cetuximab (Ctx) and anti-LDLR antibody (142F1 or 142F6) were fused to heterodimeric Fc domains respectively, with T350V/L351Y/F405A/Y407V mutations in chain A and T350V/T366L/K392L/T394W mutations in chain B^36^. To eliminate effector function for macrophage and NK cell recruitment, we introduced the L234A/L235A/P329G mutations (LALAPG)^37^ in both Fc chains. To avoid heavy and light chain mispairing, one arm was designed as single chain variable fragment (scFv) and the other in Fab format (**Fig. 2a**). We produced four Ctx-LIPTAC formats as shown in **Extended Data Fig.4a** and evaluated their degradation efficiency in Hela cells. After 24 h treatment with 50 nM of each LIPTAC, levels of EGFR were quantified by western blotting. We found that the LIPTAC with LDLR antibody as Fab and Ctx in scFv exhibited better EGFR degradation compared to the LDLR scFv format (**Extended Data Fig.4b**). LIPTAC1 containing the higher affinity Fab toward cLDLR showed the highest degradation level, suggesting that the recycling arm with higher affinity possesses better internalization efficiency.

**Fig. 2.**
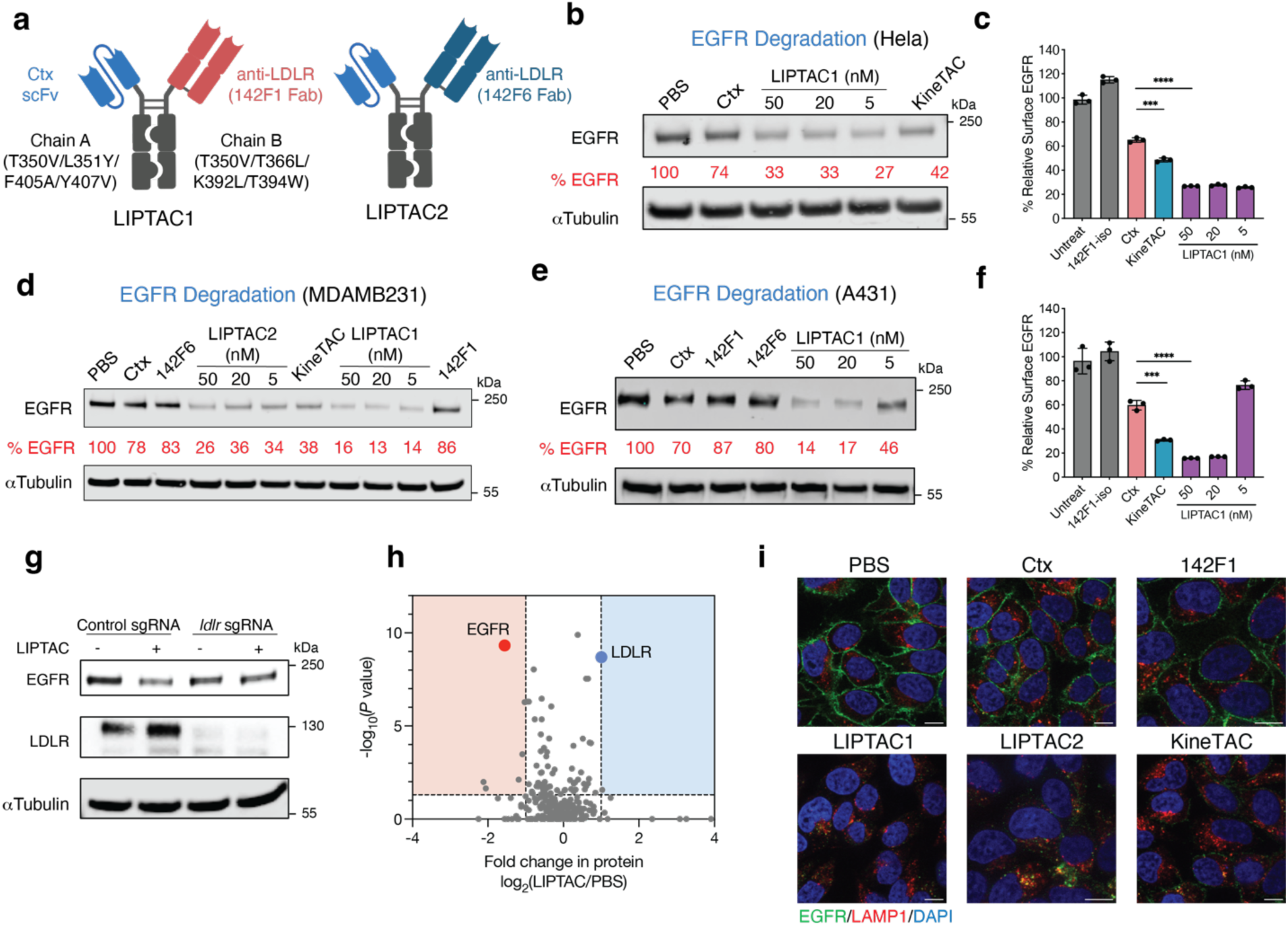
Generation of LIPTACs for degradation of EGFR. **a**, Schematic illustration of LDLR-Ctx LIPTAC bispecific constructs. **b**, Western blot showing degradation of total EGFR on HeLa cells following 24 h of treatment with Ctx-LIPTAC or 50 nM Ctx-KineTAC, and 50 nM Ctx control IgG. Data represented three biological replicates. Percent EGFR levels were quantified by ImageJ relative to PBS control. **c**, Changes in surface EGFR based on flow cytometry analysis on MDA-MB-231 cells following 24 h of 50 nM 142F1 isotype IgG, Ctx IgG, Ctx-LIPTAC, or Ctx-KineTAC treatment. Percent EGFR was determined by median fluorescence intensity (MFI) of the PE fluorescence channel of live cells. Each sample was tested in biological triplicate and error bars represented standard deviation. Statistics were calculated by unpaired two-tailed student t test. \*\*\**P* < 0.001. \*\*\*\**P* < 0.0001. **d**, Western blot showing degradation of total EGFR on MDA-MB-231 cells after 24 h treatment of 50 nM Ctx, KineTAC, monomeric LDLR isotype, or varying concetrations of LIPTACs. Data represented three biological replicates **e**, EGFR degradation in A431 cells following 24 h of Ctx-LIPTAC1 treatment. Data represented three biological replicates. **f**, Flow cytometry analysis showing degradation of surface EGFR on A431 cells following 24 h of Ctx-LIPTACs, 50 nM Ctx IgG, and 50 nM Ctx-KineTAC treatment. Each sample was tested in biological triplicate and error bars represented standard deviations. Statistics were calculated by unpaired two-tailed student t test. \*\*\**P* < 0.001. \*\*\*\**P* < 0.0001. **g**, Western blot analysis of EGFR and LDLR in LDLR knockout and control HCC1143 cells after 24 h of 5 nM LIPTAC treatment. Data represented two biological replicates. **h**, Fold-change in surface protein abundance in MDA-MB-231 cells following 48 h of treatment with or without 50 nM Ctx-LIPTAC1, as measured by quantitative proteomics analysis. N-linked cell surface glycoproteins were captured by the cell-surface capture technology^39^ and enriched by biocytin hydrazide. Surface proteins were annotated using the SURFY database^79^. **i**, Confocal microscopy images of HeLa cells treated with 50 nM of indicated bispecific antibodies or isotype controls for 24 h. Scale bar, 10 μm.

The Ctx-KineTAC utilizing the CXCL12 cytokine efficiently degrades EGFR on HeLa cells using the CXCR7 recycling receptor^19^. Thus, we compared the KineTAC and LIPTAC1. We found the LIPTAC showed somewhat higher degradation efficiency (**Fig. 2b**), indicating that LDLR might internalize and recycle faster than CXCR7. A dose-response curve for LIPTAC1 at 24h revealed that LIPTAC1 retained potent EGFR degradation activity at concentrations as low as 5 nM (**Fig. 2b**).

Given the tumor-associated nature of LDLR, we further tested degradation in multiple cancer cells. LIPTACs exhibited efficient EGFR degradation on triple-negative breast cancer cell line HCC1143 (**Extended Data Fig.4c**), the pancreatic cancer cell line PANC-1 (**Extended Data Fig.4d**), as well as the non-small cell lung cancer cell line NCI-H1975 (**Extended Data Fig.4e**). Flow cytometry demonstrated that both LIPTAC1 and the CXCL12 KineTAC efficiently degraded surface EGFR on MDA-MB-231 cells (**Fig. 2c**). Moreover, LDLR levels did not change, indicating it was not consumed in the process (**Extended Data Fig.4f**). Similar findings were observed by western blotting (**Extended Data Fig.4g**), suggesting that LDLR was recycled back. Treatment with either arm of the LIPTACs individually at 50 nM did not affect EGFR levels, indicating both targets must be brought together to cause degradation (**Fig. 2d**). Additionally, LIPTAC1 efficiently degraded EGFR with a maximal percent degradation (D_max_) of 86%, on the EGFR high expressing epidermoid carcinoma cell line A431 (**Fig. 2e**,**2f**). To assess the specificity of LDLR-mediated protein degradation, we treated LDLR knockout^38^ (KO) and control HCC1143 cells with LIPTAC1. EGFR degradation was less efficient in LDLR KO cells compared to control Cas 9 cells (**Fig. 2g**), indicating the requirement of LDLR for degradation.

To further investigate the impact of LIPTAC treatment on the proteome, we conducted quantitative mass spectrometry analysis of surface-enriched lysates^39^ following LIPTAC1 treatment in MDA-MB-231 cells. We found only a few of surface proteins showed changes in abundance (**Extended Data Fig.5**), with EGFR exhibiting the most significant reduction, supporting the high selectivity of LIPTAC1 (**Fig. 2h**). Interestingly, two tetraspanin (TSN) proteins were downregulated upon treatment (**Extended Data Fig.5**). TSN has been reported to interact with ligand-bound EGFR^40,41^, suggesting that TSN11 and TSN14 are potential neighbors of EGFR that can also be degraded. Interestingly, LDLR levels increased approximately two-fold (**Fig. 2h**), potentially due to partial competition between the 142F1 antibody and the proprotein convertase subtilisin/kexin type 9 (PCSK9) binding (**Extended Data Fig.6**), which may inhibit PCSK9-mediated LDLR degradation^42,43^. The abundance of other LDLR family members observed in our data set, including LRP1, LRP6, and LRP8, remained unchanged. This data further supports the selectivity of the LDLR antibodies.

Immunofluorescence microscopy revealed virtually complete removal of EGFR from the cell surface following 24 h of LIPTAC treatment compared to treatment with PBS, Ctx, or 142F1 clone, further highlighting that LIPTACs induce robust internalization of target proteins (**Fig. 2i**). The colocalization of lysosomal LAMP1 with intracellular EGFR further supported lysosomal shuttling by LIPTACs. Together, our findings demonstrate that LIPTAC-mediated targeted protein degradation is efficient, selective, and dependent on LDLR.

### LIPTAC-mediated degradation of multiple membrane proteins

We sought to determine whether the LIPTAC could degrade other therapeutically relevant cell surface proteins. First, we targeted PD-L1, an immune checkpoint expressed in tumor microenvironment that suppresses cytotoxic T cell function^44^. We generated a PD-L1-targeting LIPTAC by incorporating anti-PD-L1 Atezolizumab^45^ (Atz) Fab with 142F1 scFv. Atz-LIPTAC efficiently degraded PD-L1 in MDA-MB-231 cells after 24 h treatment (**Fig. 3a**, **3b**). Additionally, to determine degradation mechanisms, cells were pre-treated with Bafilomycin A (an inhibitor of lysosome acidification^46^) or MG132 (a proteasome inhibitor^47^) prior to LIPTAC1 treatment. Bafilomycin A inhibited PD-L1 degradation, while MG132 did not (Fig. 3c), suggesting that LIPTAC-mediated protein degradation occurs predominantly by delivery to lysosome.

**Fig. 3.**
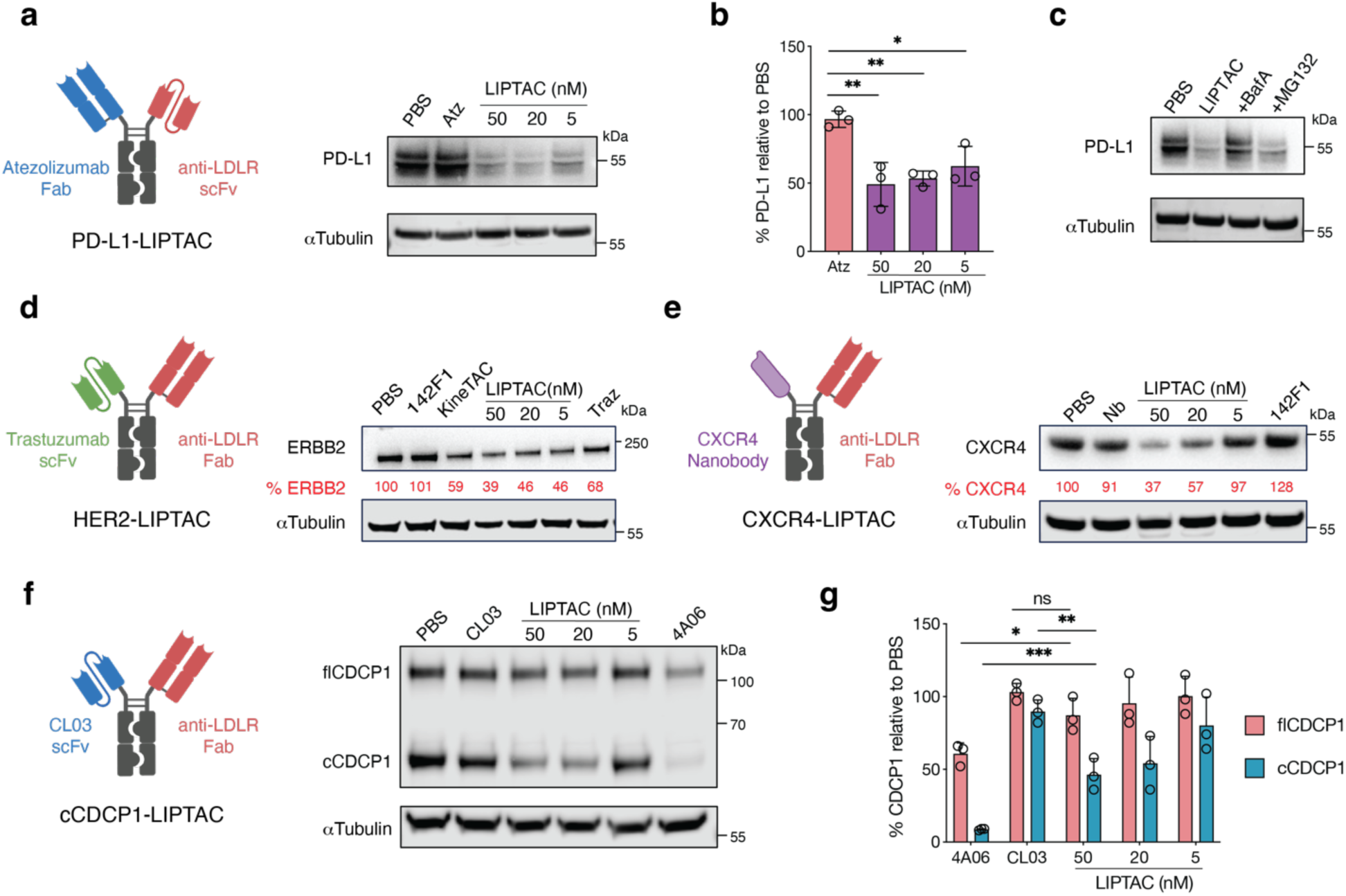
LIPTAC-mediated degradation of multiple extracellular proteins. **a**,**b**, Schematic illustration of PD-L1 targeting LIPTAC, and western blot showing total degradation of PD-L1 on MDA-MB-231 cells following 24 h treatment of 50 nM monomeric anti-PD-L1 parent, Atz, or LIPTAC containing Atz. Percent PD-L1 levels were quantified by ImageJ relative to PBS control. Each sample was tested in biological triplicate and error bars represented standard deviations. Statistics calculated by unpaired two-tailed student t test. \**P* < 0.05. \*\**P* < 0.01. **c**, Western blot analysis showing lysosome-dependent PD-L1 degradation on MDA-MB-231 cells. Cells were pretreated with either 500 nM Bafilomycin A (BafA) or 500 nM MG132 for 1 h followed by 24 h treatment with 50 nM LIPTAC. **d**, Schematic illustration of HER2-targeting LIPTAC and western blot analysis showing total HER2 degradation on MCF7 cells following 24 h treatment of monomeric Traz, LIPTAC and KineTAC. Percent ERBB2 (HER2) levels were quantified by ImageJ relative to PBS control. Data represented for at least two independent experiments. **e**, Schematic illustration of CXCR4-targeting LIPTAC and western blot analysis showing total CXCR4 degradation on HeLa cells following 24 h treatment of Nb monomer, monomeric 142F1, or LIPTAC. **f**,**g**, Schematic illustration of cleaved CDCP1-targeting LIPTAC and degradation of CDCP1 in PL45 cells following 24 h treatment of 50 nM cleaved CDCP1 (cCDCP1) binder CL03 IgG, pan-CDCP1 binder 4A06 IgG, or cCDCP1-specific LIPTAC. Each sample was tested in biological triplicate and error bars represented standard deviations. Statistics were calculated by unpaired two-tailed student t test. \**P* < 0.05. \*\**P* < 0.01. ns, not significant.

Next, we targeted human epidermal growth factor receptor 2 (HER2), which is frequently upregulated in cancer and linked to breast cancer invasiveness and tumor progression^48^. We generated a HER2-targeting LIPTAC by incorporating anti-HER2 trastuzumab (Traz) scFv with our LDLR Fab. Treatment of MCF7 cells with the Traz-LIPTAC resulted in efficient HER2 degradation (**Fig. 3d**). We then sought to evaluate degradation of multi-pass transmembrane proteins, such as G protein-coupled receptors (GPCRs). We targeted CXCR4, a chemokine receptor involved in tumor growth and metastasis^49^. The CXCR4-targeting LIPTAC was built using a CXCR4 antagonizing nanobody (Nb)^50^ on one arm and the LDLR 142F1 Fab on the other arm of the Fc. Western blotting showed significant CXCR4 degradation after 24 h treatment of Nb-LIPTAC, and not by the Nb monomer (**Fig. 3e**). The antigen-targeting arm of the LIPTAC can be flexibly incorporated into various antibody formats, including Fab fragments, scFvs, and Nbs.

Additionally, we investigated CUB domain-containing protein 1 (CDCP1), which is highly overexpressed in RAS-driven cancers and undergoes ectodomain cleavage by extracellular proteases on cancer cells but not healthy cells^51–53^. We previously developed two antibody clones: 4A06, which targets both full-length and cleaved forms of CDCP1, and CL03, which selectively recognizes the cleaved form for enhanced tumor specificity^54^. We found that the 4A06 IgG was efficiently internalized, leading to the downregulation of both full-length and cleaved CDCP1 (**Fig. 3f**). In contrast, CL03 IgG alone did not induce CDCP1 degradation. However, the CL03-based LIPTAC selectively and efficiently degraded cleaved CDCP1, without affecting the full-length form, in PL45 cells (**Fig. 3f, 3g**). Overall, these results highlight the versatility of the LIPTAC platform in selectively degrading a range of extracellular membrane proteins.

### Improved modality of target cell killing via degrader drug conjugates

Both toxin delivery by ADCs and protein degradation by eTPD require internalization, delivery, and proteolysis in the lysosome. We hypothesized that eTPD coupled to an ADC could be used to enhance the potency of an ADC in situations where the target of the ADC was not optimized for lysosomal trafficking (**Fig. 4a**). Our data here show that the Ctx-IgG only partially induces EGFR degradation (**Fig. 2e**). Given that Ctx-LIPTAC efficiently degrades EGFR, we selected Ctx ADC as a test case as a DDC.

**Fig. 4.**
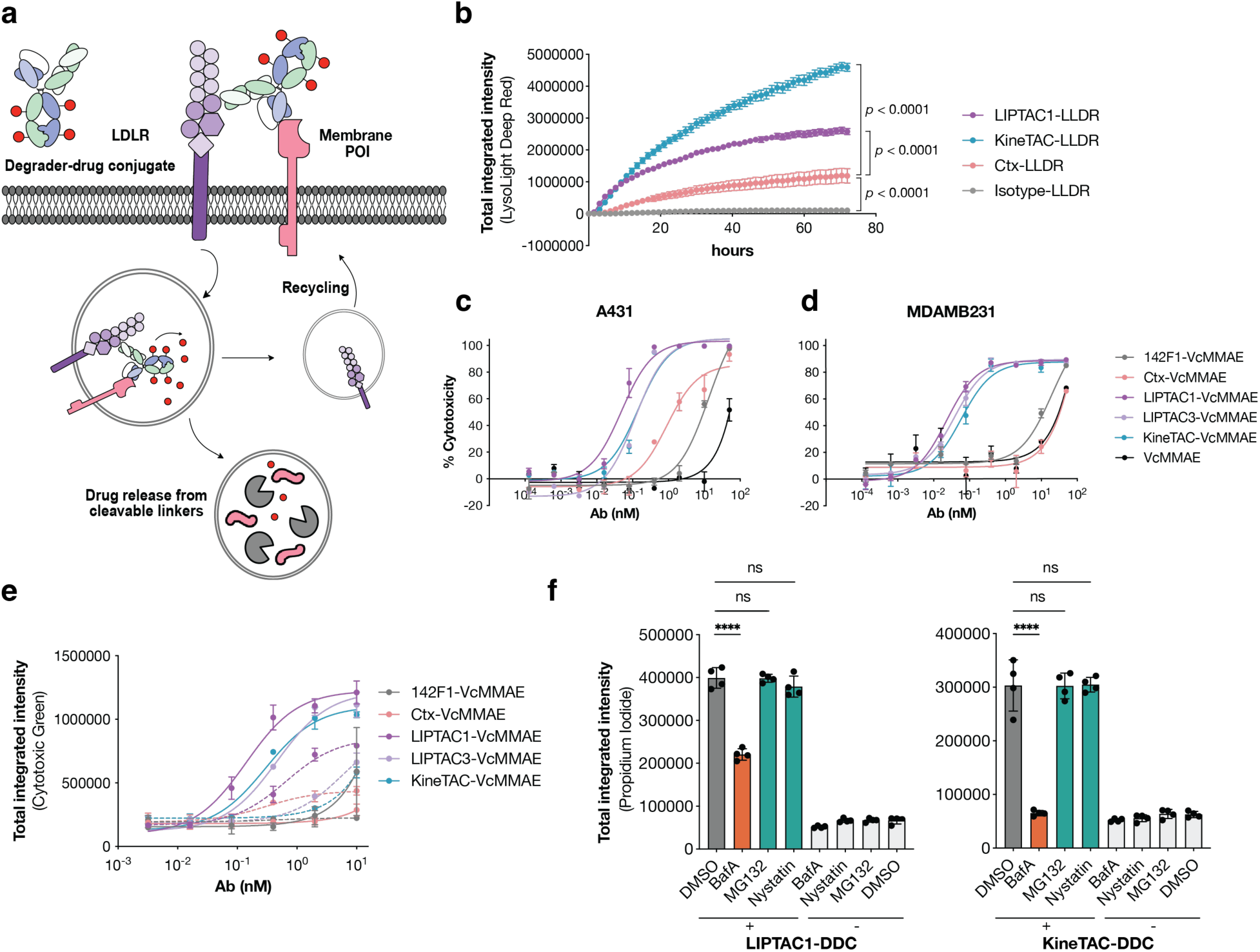
Development of degrader-drug conjugates (DDC) for potent target cell killing. **a**, Schematic illustration of LIPTAC-DDCs where the membrane POI is internalized by endocytosis and the cytotoxic payload is released from cleavable linkers in the lysosome. **b**, A payload cleavage assay in EGFR-expressing A431 cells. Cells were treated with 25 nM of antibodies conjugated with lysolight deep red (LLDR) dyes. Images were captured every 2 h for 72 h on the Incucyte. Total integrated intensity was calculated by NIRCU x μm^2^/image. Error bars represented standard deviations for four biological replicates. Statistics were calculated by one-way ANOVA and Holm-Sidak multiple comparisons test. **c**,**d**, Cytotoxicity of Ctx-ADC, monomeric 142F1-ADC, and Ctx-DDC either in LIPTAC or KineTAC formats on A431 and MDA-MB-231 cells, respectively. After 72 h incubation, cell viability was measured using the CellTiter-Glo Reagent. EC_50_ values were calculated using “One-Site Fit LogIC50” regression in GraphPad Prism 10.2. **e**, Cytotoxicity of ADCs and corresponding DDCs on A431 cells in the presence and absence of the lysosomal inhibitor BafA. Dead cells were labeled by cytotoxic green dye after 48 h of treatment. Dashed lines represented 50 nM BafA treatment together with antibodies. Total integrated intensity was calculated by GreenCU x μm^2^/image on the Incucyte. **f**, Cytotoxicity of 0.8 nM DDCs on A431 cells in the presence of 50 nM BafA, 50 nM MG132, and 1 µM Nystatin respectively. Cells treated with inhibitors alone showed no cytotoxicity at the concentrations used in DDC treatments. Dead cells were labeled by 1 µg/mL propidium iodide after 48 h of treatment. Statistics were determined by one-way ANOVA. Error bars represented standard deviations of four biological replicates. Statistics were calculated by unpaired two-tailed student t test. \**P* < 0.05. \*\*\*\**P* < 0.0001. ns, not significant.

To evaluate the drug release efficiency of DDCs, we conjugated a cathepsin-dependent fluorescent probe, LysoLight Deep Red (LLDR), to monitor the lysosomal catabolism of internalized proteins^55^. The antibody-probe conjugate was linked via a valine-citrulline (Vc) linker and remains non-fluorescent until cleavage by lysosomal cathepsins, that generates a bright fluorescent signal and serves as a surrogate for an ADC. We found that Ctx-DDC based conjugates, either in a KineTAC or LIPTAC form, showed much more efficient internalization and lysosomal trafficking compared to Ctx-IgG ADC (Fig. 4b, **Extended Data Fig.7a**), suggesting enhanced drug delivery efficiency for the DDC.

Next, we evaluated the toxicity of Ctx-DDC by conjugating them to VcMMAE as the cytotoxic payload. MMAE is an auristatin derivative that inhibits tubulin polymerization with a cleavable VC linker^56^. To compare the cell killing potency of DDCs and Ctx ADC, we conjugated VcMMAE to monomeric 142F1, LIPTAC1, LIPTAC3, KineTAC, and Ctx IgG respectively. The average drug–antibody ratio (DAR) was 2.6, as estimated using 2,4-dinitrophenol (DNP)-PEG4 conjugation with the same method (**Extended Table 1**). First, we confirmed that antibodies after drug conjugation retained binding to EGFR^+^ cells (**Extended Data Fig.7b**). Next, we evaluated the potencies of the DDCs in EGFR^high^ A431 cells. The Ctx ADC exhibited an IC_50_ of 0.9 nM, while the LIPTAC and KineTAC DDCs significantly enhanced the potency by 18-fold and 7-fold, respectively (**Fig. 4c**). We then assessed the cytotoxicity in EGFR^medium^ MDA-MB-231 cells, which others have shown are relatively insensitive to Ctx-VcMMAE due to the low receptor expression^57^. After 72 h of treatment, the Ctx ADC induced only moderate cytotoxicity, even at the highest ADC concentration (**Fig. 4d**). The single arm anti-LDLR 142F1-VcMMAE also exhibited cell killing weak potency, suggesting that LIPTAC binding is primarily driven by the higher-affinity Ctx arm (**Fig. 4d**, **Extended Data Fig.7b**). Remarkably, all bispecific DDCs, including LIPTAC1-VcMMAE, LIPTAC3-VcMMAE, and KineTAC-VcMMAE, potently killed the target cells (**Fig. 4d**), with IC_50_ values of 23 pM, 38 pM, and 59 pM, respectively.

We observed similar DDC-mediated killing potency on PANC-1 cells which expressed similar levels of EGFR as MDA-MB-231 (**Extended Data Fig.7c**). None of the antibodies without payload were significantly toxic to cells over three days of treatment (**Extended Data Fig.7d**), suggesting that the cytotoxicity is mediated by internalization and release of the degrader conjugated MMAE. The cytotoxicity of ADC and DDCs was minimal on EGFR negative MCF7 cells (**Extended Data Fig.7e**).

We also evaluated the lysosomal dependency of cell killing using an incucyte-based killing assay. Accumulation of cell death was still detectable when cells were treated with as low as 0.4 nM of DDCs, while the cell death was not detectable with Ctx-ADC or 142F1-ADC (**Fig. 4e**, **Extended Data Fig.7f**). Bafilomycin A treatment abolished the cytotoxicity of both ADC and DDCs, further indicating the requirement of lysosomal release of cytotoxic payload (**Fig. 4e**). We further treated cells with chemical inhibitors of several endocytic pathways. Inhibitors of proteasomes (MG132) or caveolar-mediated endocytosis (Nystatin)^58^ did not significantly affect the cytotoxicity of the DDCs, highlighting receptor-mediated lysosomal delivery (**Fig. 4f**). Together, these data suggest that DDCs improve lysosomal delivery of cytotoxic payload and enhance the cytotoxic potency.

### Degrader-drug conjugates improved cytotoxicity on cells with moderate antigen expression levels

To evaluate whether DDCs can enhance the drug delivery efficiency of these ADCs, we selected four clinically relevant ADC targets: anti-B cell maturation antigen (BCMA) Belantamab^59,60^, anti-CD19 Loncastuximab^61^, anti-trophoblast cell surface antigen 2 (TROP2) Sacituzumab^62^, and anti-HER2 Trastuzumab^63^. We engineered these therapeutically relevant antibodies into corresponding LIPTAC formats and conjugated them to VcMMAE as similar DARs to EGFR-targeting DDCs.

BCMA is highly expressed on malignant plasma cells and represents a validated target in multiple myeloma. Belantamab mafodotin (Blenrep) is a novel ADC composed of an anti-BCMA antibody conjugated to MMAF via a non-cleavable linker^64^. Our Belantamab-LIPTAC demonstrated more efficient internalization than Belantamab in RPMI-8226 cells (**Fig. 5a**). Both the conjugated ADCs and DDCs retained binding affinity comparable to their unconjugated counterparts (**Extended Data Fig.8a**). In the luciferase-based viability assay, LIPTAC DDCs improved potency in RPMI-8226 cells by 4-fold, reducing the EC₅₀ from 85 nM to 22 nM. The effect was not observed in MM1.S cells with higher BCMA expression (**Extended Data Fig.8b**). Notably, by Annexin V and propidium iodide staining, LIPTAC DDCs significantly enhanced cell death by 18-fold in RPMI-8226 and 5-fold in MM1.S cells (**Fig. 5b**, **Extended Data Fig.8c**). This difference may be due to incomplete killing by DDCs in BCMA^+^ cells, with Annexin V staining capturing early apoptotic events that are not detected by viability assays. Overexpression of BCMA in RPMI-8226 cells rendered both LIPTAC DDC and Belantamab ADC comparably potent (**Extended Data Fig.8d**, **8e**), suggesting that LIPTAC-mediated trafficking offers the greatest advantage in cells with moderate antigen expression.

**Fig. 5.**
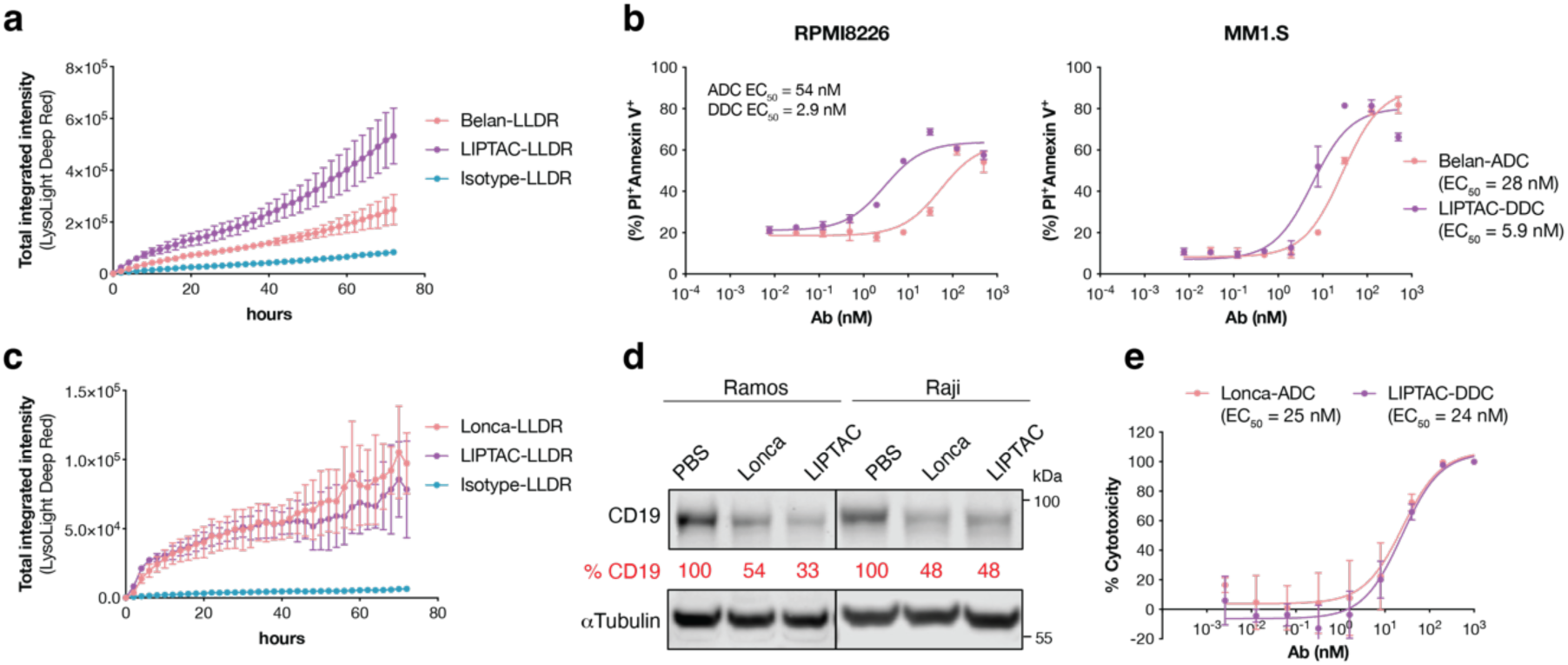
Comparison of clinically approved ADCs and the DDC counterparts. **a**, Antibody internalization assay in RPMI8226 cells, a multiple myeloma cell line, treated with 100 nM of LLDR dye-labeled antibodies. An anti-SARS-CoV-2 spike antibody CC12.1 was used as the isotype control^80^. Images were captured every 2 h for 72 h on the Incucyte. Each sample was tested in biological triplicate, and error bars represent the standard deviations. **b**, Cytotoxicity of anti-BCMA Belantamab (Belan) ADC or the corresponding LIPTAC-DDC after 4 days incubation with RPMI8226 and MM1.S cells. To monitor cell death, cells were stained with APC-annexin V and propidium iodide (PI) and analyzed by flow cytometry. Each sample was tested in biological triplicate and error bars represent standard deviations. **c**, Antibody internalization assay in Raji cells, a B-cell lymphoma cell line, treated with 100 nM of LLDR dye-labeled antibodies. Images were captured every 2 h for 72 h on the Incucyte. Each sample was tested in biological triplicate and error bars represented the standard deviations. **d**, Western blot analysis showing total degradation of CD19 on Ramos and Raji cells after 24 h treatment of 50 nM antibodies. Percent CD19 levels were quantified by ImageJ relative to PBS control. **e**, Cytotoxicity of an anti-CD19 Loncastuximab (Lonca) ADC or LIPTAC-DDC after 4 days incubation with Ramos cells. Cell viability was measured using the CellTiter-Glo Reagent. Each sample was tested in biological triplicate and error bars represented standard deviations. EC_50_ values were calculated using “One-Site Fit LogIC50” regression in GraphPad Prism 10.2.

Next, we investigated CD19, a protein strictly and ubiquitously expressed on B cells across multiple developmental stages. Loncastuximab tesirine is an anti-CD19 ADC where the payload is a PBD dimer that inhibits DNA metabolic processes via a cleavable dipeptide linker^61^. CD19 has rapid internalization kinetics, with colocalization of anti-CD19 antibodies and lysosomal compartments observed as early as 15 minutes post-treatment^65^. CD19-LIPTAC did not enhance internalization (Fig.5c, **Extended Data Fig.9a**). Although the LIPTAC improved CD19 degradation, Loncastuximab (Lonca) itself efficiently internalized and degraded CD19 (Fig. 5d). Consequently, the cytotoxicity of the LIPTAC DDC was comparable to Lonca-ADC (Fig. 5e, **Extended Data Fig.9b**,**9c**), indicating minimal benefit from the LIPTAC format in this context.

Similarly, Trastuzumab is known to induce HER2 internalization and downregulation upon binding^66,67^. Although HER2-LIPTAC modestly improved internalization (**Extended Data Fig.10a**), it did not enhance the cytotoxicity in HER2^high^ BT474 cells or HER2^low^ T47D cells (**Extended Data Fig.10b**,**10c**). A similar outcome was observed for TROP2-targeting Sacituzumab ADCs. TROP2 undergoes rapid internalization upon antibody binding, with a reported internalization half-life (t₁/₂) of approximately 30 minutes^68^. Despite moderate improvements in internalization with TROP2-LIPTAC (**Extended Data Fig.10d**), no significant increase in cytotoxicity was observed in TROP2^high^ A431 or TROP2^medium^ PL45 cells (**Extended Data Fig.10e-10g**). Interestingly, LIPTAC DDC did show greater potency compared to monomeric Sacituzumab ADC (**Extended Data Fig.10e-10g**), suggesting that increased avidity or receptor clustering could contribute to improved lysosomal delivery for the ADC.

Overall, these results indicate that LIPTAC-based DDCs can enhance cytotoxicity in specific contexts, particularly in target-low settings or when the parental ADC is limited by suboptimal internalization. When the ADC is in high abundance and a good recycler the benefit of a DDC is clearly less. These findings underscore the importance of target selection and receptor biology in guiding DDC design.

## Discussion

TPD has emerged as an exciting new modality for potent degradation of functionally important proteins inside and outside the cell. Whereas iTPD has principally focused on two intracellular E3 ligases, due to limitations of chemical materials that can bind cereblon and VHL, eTPD can readily access scores of extracellular recycling receptors with routine antibody generation and established bispecific constructs or conjugations. Thus, eTPD allows for greater tissue selectivity by proper choice of the degrader system that matches the expression of the POI on disease versus healthy tissues. The LDLR-based LIPTAC expands opportunities for eTPD. LDLR is an attractive recruiter for catalytic and durable degradation due to its rapid recycling kinetics. It can complete one endocytic cycle approximately every 12 minutes and recycle up to 150 times^69^. Here, we show that LIPTAC enables efficient, selective, and LDLR-dependent lysosomal degradation of a diverse range of therapeutically relevant membrane proteins, including both single-and multi-pass transmembrane proteins, as well as the cleaved, tumor-specific neoepitopes such as CDCP1.

We further demonstrate that eTPD technologies such as LIPTAC and KineTAC can complement ADCs by improving intracellular trafficking and lysosomal delivery, which are critical determinants of ADC efficacy^70^. Recent biparatopic and bispecific ADC designs (e.g., HER2×CD63^10^, HER2×PRLR^71^, and TROP2×APLP2^11^) have leveraged receptor clustering and dual-antigen engagement to enhance lysosomal targeting^9^. Building on this, we show that antibody-based degraders can similarly drive efficient endolysosomal delivery and serve as potent payload vehicles. In head-to-head comparisons, EGFR-targeting Ctx-DDCs outperformed Ctx-based ADCs in lysosomal trafficking and cytotoxicity. Tumor selectivity was conferred by the high-affinity Ctx arm^72^, as LDLR-targeting alone exhibited weak activity. LIPTAC-DDC exhibited minimal toxicity to cells without EGFR expression, further highlighting that LIPTAC specificity stems from the tumor antigen-binding domain. The bispecific nature that binds the POI and the degrader system can allow for greater tumor specificity by choosing degrader systems matched to the tumor and not healthy cells. The LDLR is significantly upregulated in many cancer cells relative to normal cells providing an advantage in this regard.

The therapeutic potency of ADCs depends on antigen abundance, receptor internalization, and payload potency. Our results suggest that LIPTACs can enhance payload delivery particularly in contexts where conventional ADCs are limited, such as receptors with moderate expression or poor internalization (e.g., BCMA on RPMI-8226 cells). However, this advantage diminished in BCMA-overexpressing cells and was not observed for efficiently internalizing targets like CD19 or TROP2, where LIPTAC and KineTAC formats performed comparably to approved ADCs (e.g., Loncastuximab, Sacituzumab). Reduced avidity and lower binding affinities of LIPTACs also likely contributed to these differences.

The DDC format may expand druggability to challenging targets with poor internalization kinetics or low surface abundance, such as post-translational modifications^73,74^, tumor-specific neo-antigens^75^, or glycosylphosphatidylinositol (GPI)-anchored proteins^76^. Future optimization of antigen-targeting arms and use of heterobifunctional degraders like LIPTAC could reduce DAR requirements, minimize off-target effects, and broaden the therapeutic window.

Finally, a major limitation of eTPD approaches is incomplete degradation, which may be insufficient to fully suppress oncogenic signaling. The DDC format overcomes this by delivering cytotoxic payloads directly to tumor cells, independent of full protein clearance. This modular platform can be expanded to deliver diverse therapeutic cargos, including radionuclides, kinase inhibitors, or proteolysis-targeting chimeras (PROTACs). In summary, the DDC modality represents a promising next-generation hybrid strategy for targeted cancer therapy.

## Acknowledgement

We thank Dr. Brandon Holmes, Dr. Jon Ostrem and Dr. Rohit Bhadoria for their assistance with discussions, and the Wells Lab broadly for helpful discussions and expertise. We thank Paul Burroughs for providing the BCMA overexpressing RPMI8226 cells. We are grateful to generous support from NIH-1R01CA248323-01(J.A.W), NIH-R35GM122451 (J.A.W.), the Hind Professorship in Pharmaceutical Sciences (J.A.W), and R01CA276207 (J.A.O.). K.S. is supported by a Helen Hay Whitney Foundation Fellowship. Z.Y. is supported by a National Institute of General Medical Sciences F32 Postdoctoral Fellowship. K.K. is supported by a graduate fellowship funded by the National Science Foundation.

## Author Contributions

F.Z. and J.A.W. conceived and designed the study. F.Z. performed phage display, antibody screening and characterization experiments. F.Z., Y.W., Y.Z., S.G. and K.K. cloned and expressed the recombinant proteins. F.Z. and Z.Y. performed the western blotting experiments. F.Z., Y.W., and K.K. labeled the antibodies with dyes or payloads and performed the internalization and cytotoxicity assays. K.S. and T.M.P.C. performed the proteomics and processed the data. K.M. provided the confocal microscopy data. A.I and J.A.O. provided the LDLR KO cells. F.Z. and J.A.W. wrote the manuscript and all authors reviewed and edited the manuscript.

## Materials and Methods

### Plasmid construction

All the IgGs were constructed in a pcDNA3.4 vector that expresses the light chain and heavy chain, respectively for mammalian expression. For generating bispecific antibody, the heavy chain variable regions or scFvs were cloned into zymework-A mutant Fc and zymework-B Fc sequences respectively. Antigen was cloned into pFuse vector with IL-2 signal peptide followed by tobacco etch virus (TEV) protease, Fc, and Avitag sequence in the C-terminus. All the Fabs were constructed in a dual-expression pBL347 vector that expresses the light chain and the heavy chain with the pelB and the stII signal peptides, respectively, for the periplasm expression.

### Cell lines

HEK293T, HeLa, MDA-MB-231, PANC-1 and A431 were cultured in DMEM (ThermoFisher Scientific) with 10% fetal bovine serum (FBS) and 1% penicillin– streptomycin. MCF7 cells were cultured in DMEM with 10% FBS, 1% penicillin-streptomycin, 1% sodium pyruvate, and 1% non-essential amino acid. NCI-H1975, HCC1143, PL45, BT474, Raji, Ramos, RPMI8226, and MM1.S cells were cultured in RPMI-1640 (ThermoFisher Scientific) with 10% FBS and 1% penicillin–streptomycin. HCC1143 LDLR KO cells^38^ were provided by Dr. James Olzmann and were supplemented with 100 μg/mL hygromycin (Sigma-Aldrich). of Expi293F cells were cultured in FreeStyle 293F medium.

### Phage display

Phage selection was done as described previously^77^. In brief, library E and UCSF library were incubated with streptavidin-coated magnetic beads pre-conjugated with biotinylated Fc protein to remove nonspecific binders. Unbounded phages were then incubated with streptavidin-coated magnetic beads pre-conjugated with biotinylated cLDLR-TEV-Fc antigens. After 4 washes, antigen-bound phages were eluted from beads by incubating with 1 μM TEV protease for 20 min. In total, four rounds of selections were performed with a decreasing concentration of cLDLR-antigen (1000, 50, 20, 10 nM). From round 3, the phage library was first enriched by protein A magnetic beads to deplete nondisplayed or truncated Fab phage before each round of the selection.

### Phage ELISA

384-well Maxisorp plates were coated with Neutravidin (10 μg/mL) overnight at 4 °C and subsequently blocked with BSA (2% w/v) for 1 h at RT. 20 nM biotinylated cLDLR antigens were captured on the NeutrAvidin-coated wells for 30 min followed by the addition of 1:5 diluted single-colony phage for 1 h. The secondary antibodies were either a horseradish peroxidase (HRP)-conjugated anti-M13 phage antibody (Sino Biological) for phage ELISA or an anti-human IgG antibody (Sigma-Aldrich) for recombinant protein ELISA. The ELISA plates were washed three times after each incubation, and antibody binding was detected by TMB substrate (VWR) and read at 450 nm.

### Protein expression

IgGs and antigen were expressed in Expi293F cells in a 30 mL scale. In brief, 24 μg of DNA was added to 3 mL of OptiMEM, followed by 24 μL of FectoPro transfection reagents. After 10 min of incubation, 27 mL of Expi293F cells at 3 millions/mL were added and shake at 37°C. On the second day, 300 μL of 300 mM vaporic acid and 270 μL of 45% glucose were added to the cells. After 5 days of transfection, cells were harvested, spun down at 4,000 g for 20 min and filtered by 0.45 μm steri-flip. Supernatants were then incubated with Sepharose A resin for 2 h, proteins were then eluted by 0.1 M acetic acid and neutralized by Tris pH 11. Proteins were buffer changed 3 times in PBS in amicon tubes. Fabs were expressed in *Escherichia coli* C43 (DE3) Pro+ grown in an optimized TB autoinduction medium at 37 °C for 6 h, cooled to 30 °C for 18 h. Cells were harvested by centrifugation and lysed using B-PER lysis buffer. The lysate was incubated at 60 °C for 20 min and centrifuged to remove the inclusion body. The Fabs were purified by Sepharose A resin via affinity chromatography and buffer exchanged in PBS for further characterization. Purity and integrity of all proteins were assessed by SDS–PAGE.

### Recombinant protein ELISA

384-well Maxisorp plates were coated with Neutravidin (10 μg/mL) or anti-histag antibody (Invitrogen, 2 μg/mL) overnight at 4 °C and subsequently blocked with BSA (2% w/v) for 1 h at RT. 20 nM of antigens were captured onto pre-coated wells for 1h. Recombinant full-length LDLR, LRP2, LRP8, and VLDLR proteins were purchased from ACROBiosystems. For polyspecificity ELISA, autoantigens cardiolipin (Sigma, 50 μg/mL), insulin (Sigma, 1 μg/mL), lipopolysaccharid (LPS, InvivoGen, 10 μg/mL), and single-stranded DNA (ssDNA, Sigma, 1 μg/mL), were directly coated onto plates overnight 4 °C. After three times of wash, serially diluted Fabs were added to the plates and incubated for 1 h at RT. After three times of wash, 1:5000 diluted peroxidase-anti-human IgG (H+L) (Jackson ImmunoResearch) were added to the plates and incubated for 30 min. After three times of wash, antibody binding was detected by TMB substrate (VWR), quenched by 1 M phosphoric acid, and read at 450 nm.

### Flow cytometry

Cells were collected by centrifugation at 400 g for 5 min. Pellets were washed once with PBS + 1% BSA. Cells were incubated with fluorophore-conjugated antibodies in PBS + 1% BSA for 15 min at RT or 30 min at 4 °C. Cells were washed three times and resuspended in cold PBS for flow analysis. Antibodies used included APC anti-LDLR (Invitrogen, Cat# MA5-40994, 1:400), PE anti-human EGFR (Invitrogen, Cat#MA5-28544,1:400), APC anti-human CXCR4 (Biolegend, Cat#306509, 1:400), Alexa fluor 647 goat anti-human IgG (H+L) (Invitrogen, Cat#A-21445, 1:1000), APC Annexin V (BioLegend, Cat#640920, 1:500), Alexa fluor 647-conjugated protein A (Invitrogen, Cat# P21462, 1:1000). Dead cell staining included propidium Iodide (Biolegend, Cat#421301, 1:250), and LIVE/DEAD™ fixable violet dead cell stain kit (Invitrogen, Cat# L34964). Flow cytometry was performed using a CytoFLEX cytometer (Beckman Coulter, v.2.3.1.22) and CytoExpert software (v.2.3.1.22). Data were analyzed with FlowJo (v.10.8.0).

### Biolayer interferometry

BLI experiments were performed at room temperature using an Octet RED384 instrument (ForteBio). 20 nM biotinylated antigens were immobilized to an optically transparent SA biosensor (ForteBio). Different concentrations of antibodies in kinetics buffer (PBS, 0.05% Tween-20, 0.2% BSA) were used as the analyte in a 384-well microplate (Greiner Bio-One). Affinities (K_D_s) were calculated by a global fit analysis and by a 2:1 heterogeneous ligand model using the Octet RED384 Data Analysis HT software.

### Epitope binning by BLI

Anti-LDLR antibodies were binned into epitope specificities using an Octet RED384 system. 20 nM of biotinylated cLDLR-Fc antigens were captured using streptavidin biosensors (Fortebio). After antigen loading, a saturating concentration of antibodies (200 nM) was added for 10 min. Competing concentrations of antibodies (40 nM) were then added for 5 min to measure binding in the presence of saturating antibodies. All incubation steps were performed in PBS/0.05% Tween-20/0.2% BSA. For PCSK9 epitope binning, all incubation steps and protein dilution were performed at acidic endosomal pH to increase PCSK9 binding affinity^78^. 200 nM PCSK9 D374Y (AcroBiosystems) was added for 10 min after antigen loading. Then 40 nM of PCSK9 D374Y, 142F1, or 142F6 was added for 5 min.

### LDL uptake assay

LDL uptake was measured by the Image-iT™ pHrodo™ Red Low Density Lipoprotein Uptake Kit (Thermo Scientific, Cat#I34360) following manufacturer’s instructions. Briefly, HeLa cells were seeded at 5000 cells/well on a 96-well polystyrene tissue culture treated plate (Corning, Cat#3596). The next day, media was removed and cells were serum starved for 12 h. Next, cells were pretreated with unlabeled LDL, heparin, and anti-LDLR Fabs for 30 min at 37 °C respectively, followed by pHrodo™ red-labeled LDL treatment. Cells were then imaged by Incucyte (Sartorius) every 1 to 2 h. Internalization was calculated by total integrated intensity (ROCU x μm^2^/image) on the Incucyte software.

### Degradation experiments

Cells were plated in 6-or 12-well plates and grown to ∼70% confluency before treatment. On the next day, cell culture medium was aspirated, various concentrations of antibodies in 1 mL of culture medium were then added to each well. Cells were incubated for 24 h at 37 °C for flow cytometry or western blotting experiments.

### Western blotting

Cells were lifted with PBS+0.05% EDTA, transferred to Eppendorf tubes, spun down at 500 g for 4 min, and wash 2 times with PBS. Then cells were lysed with 1× RIPA lysis buffer (EMD Millipore) with cOmplete mini protease inhibitor cocktail (Sigma-Aldrich) at 4 °C for 20 min. Lysates were centrifuged at 20,000g for 10 min at 4 °C. Protein amounts were quantified by Rapid Gold BCA Protein Assay Kit (Pierce). Lysates were mixed with 4× Nupage LDS Sample Buffer (Invitrogen) and 2-mercaptoethanol, and then run on NuPAGE™ 4-12% Bis Tris Protein Gels (Thermo Fisher Scientific). Proteins were transferred to polyvinylidene difluoride membranes using the iBlot2 Western Blotting Transfer System (Thermo Scientific). Membranes were blocked with TBS + 5% BSA + 0.5% Tween for 1h, and stained with primary antibodies overnight. After three times of washing, membranes were stained with secondary antibodies for 1 h at RT. After three times of washing, membranes were imaged with a LICOR imager or the ChemiDoc MP imaging system (BioRad). Antibodies used included rabbit anti-human EGFR (Cell Signaling Technology, Cat#4267S, 1:1000), rabbit anti-human PD-L1 (Cell Signaling Technology, Cat#13684S, 1:1000), rabbit anti-human CXCR4 (Cell Signaling Technology, Cat#64837S, 1:1000), rabbit anti-human ERBB2 (Cell Signaling Technology, Cat#4290S, 1:1000), rabbit anti-human CDCP1 (Cell Signaling Technology, Cat#13794S, 1:1000), rabbit anti-human CD19 (Cell Signaling Technology, Cat# 90176T, 1:1000), mouse anti-human β-tubulin (Cell Signaling Technology, 3873S, 1:3000), goat anti-human LDLR (R&D Systems, Cat#AF2148, 1:1000), IRDye 800CW goat anti-rabbit IgG (LI-COR Biosciences, Cat# 926-32211), IRDye 680RD goat anti-mouse IgG (LI-COR Biosciences, Cat# 926-68070, 1:5000), IRDye 800CW donkey anti-goat IgG (LI-COR Biosciences, Cat# 926-32214, 1:5000), peroxidase goat anti-rabbit IgG (H+L) (Jackson ImmunoResearch, Cat# 111-035-144, 1:5000).

### Confocal microscopy

HeLa cells were plated on the chambered coverslip (Ibidi, 8-well uncoated) and incubated for 24 h at 37°C. Cells were then treated with 50 nM bispecific or control antibodies in complete growth medium. After 24 h of incubation at 37°C, medium was aspirated, and cells were washed with PBS. Cells on the coverslips were fixed with paraformaldehyde (PFA) for 15min at RT, then permeabilized with 0.1% Triton-X in PBS for 10 min at RT. After washed 3 times by PBS, the resulting sample were stained with anti-LAMP1 rabbit antibody (Cell Signaling Technology, Cat# 9091T), anti-EGFR mouse antibody (Thermo Scientific, Cat#MA5-13070), and DAPI (Cell Signaling Technologies). Goat anti-rabbit IgG 488 (Invitrogen, Cat#A-11008) and goat anti-mouse IgG 647 (Invitrogen, Cat#A-21240) were stained for visualization. Samples were imaged using a Nikon Ti Microscope with a Yokogawa CSU-22 spinning disk confocal and a 60x objective lens; 405-, 488-and 647-nm lasers were used to image DAPI, LAMP1 and EGFR, respectively. Images were deconvoluted and processed using NIS-Element (v5.21.03) and Fiji software (v2.1.0) packages.

### Cell culture/stable isotope labeling using amino acids in cell culture (SILAC) labeling

MDA-MB-231 cells were grown in DMEM for SILAC (Thermo Fisher) with 10% dialyzed FBS (Gemini). Medium was also supplemented with either light L-[^12^C_6_,^14^N_2_]-lysine/l- [^12^C_6_,^14^N_4_]-arginine (Sigma) or heavy L-[^13^C_6_,^15^N_2_]-lysine/L-[^13^C_6_,^15^N_4_]-arginine (Cambridge Isotope Laboratories). Cells were maintained in SILAC medium for five passages to ensure complete isotopic labeling. Heavy-labeled cells were treated with PBS control and light-labeled cells were treated with 50 nM bispecific LIPTAC for 48 h before cells were collected. Cells were then used to prepare surface-proteome enrichment.

### Mass spectrometry

For proteomic analysis, cells were processed following established cell surface capture methods^39^. Approximately 2 million SILAC-labeled cells were first washed in PBS (pH 6.5) before the glycoproteins were oxidized with 1.6 mM sodium periodate (Sigma) in PBS (pH 6.5) for 20 min at 4 °C. Cells were then biotinylated via the oxidized vicinal diols with 1 mM biocytin hydrazide (Biotium) in the presence of 10 mM aniline (Sigma) in PBS (pH 6.5) for 90 min at 4 °C. Cell pellets were lysed with a 2× dilution of commercial RIPA buffer (Millipore) supplemented with 1× protease inhibitor cocktail (Sigma) and 2 mM EDTA (Sigma) for 10 min at 4 °C. Cells were further disrupted with probe sonication (20% amplitude, 5 min, 4 °C), followed by cell debris removal (20,000xg, 10 min, 4 °C), and the clarified cell lysates were then incubated with 50 µL of high-capacity NeutrAvidin-coated agarose beads (Thermo) in Poly-Prep chromatography columns (Bio-Rad) for 2 h at 4 °C to isolate biotinylated glycoproteins. To enrich for biotinylated proteins, the resin was washed sequentially with 5 mL of 1× RIPA (Millipore) plus 1 mM EDTA, 5 mL high-salt PBS (20 mM phosphate (pH 7.4) with 1 M NaCl (Sigma)) and 5 mL of denaturing urea buffer (50 mM ammonium bicarbonate and 2 M urea). All wash buffers were heated to 42 °C before use. Proteins on the beads were next reduced, carbidomethylated, digested and desalted using the Preomics iST mass spectrometry sample preparation kit (Preomics) per the manufacturer’s recommendations. After desalting, samples were dried, resuspended in 0.1% formic acid and quantified using the Pierce peptide quantification kit (Thermo Scientific) before liquid chromatography–tandem mass spectrometry analysis. Liquid chromatography–tandem mass spectrometry was performed using a Bruker NanoElute chromatography system coupled to a Bruker timsTOF Pro mass spectrometer. Peptides were separated using a prepacked IonOpticks Aurora (25 cm × 75 μm) C18 reversed-phase column (1.6-µm pore size, Thermo) fitted with a CaptiveSpray emitter for the timsTOF Pro CaptiveSpray source. For all samples, 200 ng of resuspended peptides was injected and separated using a linear gradient of 2–23% solvent B (solvent A: 0.1% formic acid and 2% acetonitrile; solvent B: acetonitrile with 0.1% formic acid) over 90 min at 400 µl min–1 with a final ramp to 34% B over 10 min. Separations were performed at a column temperature of 50 °C. Data-dependent acquisition was performed using a timsTOF PASEF tandem mass spectrometry method (TIMS mobility scan range of 0.70–1.50 V•s cm–2, mass scan range of 100–1,700 m/z, ramp time of 100 ms, 10 PASEF scans per 1.17 s, active exclusion of 24 s, charge range of 0–5 and minimum MS1 intensity of 500). The normalized collision energy was set at 20.

### Mass spectrometry data analysis

LC-MS-MS data was analyzed using PEAKS online Xpro 1.6 (Bioinformatics Solutions Inc.; Ontario, Canada). Spectral searches were performed in PEAKS Q (de novo assisted Quantification) mode. The precursor mass error tolerance was set to 20 ppm, and the fragment mass error tolerance was set to 0.5. Peptides containing 6 and 45 amino acids in length were then searched in a semi-specific trypsin/LysC digest mode against a proteome file that contains human cell surface proteins^79^. Carbidomethylation (+57.0214 Da) on cysteines was a set static modification; methionine oxidation (+15.994), and the isotopic labels (13C(6)15N(2); 13C(6)15N(4)) were set variable modifications. Quantified peptides were matched between three experimental replicates and peptide enrichments were normalized based on the total ion chromatograph (TIC). SILAC-labeled protein ratios were further analyzed if the proteins were identified by more than one peptide and present in at least two experimental replicates. A p-value of 0.05 and two-fold protein ratio differences were set as cut-offs to determine if protein abundance differences were significant between vehicle treated cells and LIPTAC-treated cells. Proteomic data is available on PRIDE with PXD064642 accession code.

### Lysolight deep red assay

Antibodies were labeled using the LysoLight Antibody Labeling Kits (Invitrogen, Cat# L36003) following manufacturer’s instructions. Briefly, antibodies were labeled with LLDR with a molar ratio of 1:6 in the presence of 100 mM sodium bicarbonate (pH 8.4) for 2 h at RT. Antibodies were then purified with 7k Zeba dye and biotin removal columns (Thermo Scientific, Cat#A44297). Cells were seeded at 5000/well on a 96-well polystyrene tissue culture treated plate (Corning, Cat# 3596). The next day, media was removed, treated with LLDR-labeled antibodies, and then imaged on the Incucyte every 2 h for 72 h.

### *In vitro* ADC assays

IgGs were labeled with NHS ester-PEG4-ValCit-PAB-MMAE (BroadPharm, Cat#BP-25503) with 1:6 or 1:10 molar ratio at RT for 2 h with 100 mM sodium bicarbonate. Antibodies were then desalted using the Pierce Zeba desalt spin columns (Thermo Scientific). For adherent cells, 5000 cells/well were seeded on a 96-well polylysine-coated white plate (Corning, Cat#3917). The next day, media was aspirated, MMAE-labeled IgGs were added and incubated for 72 h. For suspension cells, 6000 cells/well were incubated with ADCs or DDCs for 96 hours. Viability was measured using CellTiter-Glo Reagent (Promega). For incucyte-based killing assay, the cells were treated with ADCs or DDCs, cytotoxic green dye (Sartorius, Cat#4633, 1:10000) or propidium Iodide (Biolegend, Cat#42130, 2 µg/mL). For compound inhibitor assay, 0.8 nM of DDCs were incubated with 1 µg/mL of propidium Iodide with or without 50 nM Bafilomycin A (Santa Cruz Biotechnology, Cat#sc-201550A), 50 nM MG132 (Selleck Chemicals, Cat#S2619), or 1 µM Nystatin (MedChem Express, Cat# HY-17409), respectively.

### Characterization of DAR

IgGs were side-by-side labeled with DNP-PEG4-NHS ester (MedChem Express, Cat# HY-140614) with the same molar ratio as MMAE conjugation, and incubated at RT for 2 h with 100 mM sodium bicarbonate. Antibodies were then desalted using the Pierce Zeba desalt spin columns (Thermo Scientific). The absorbance of conjugated antibodies at 280 nm and 360 nm was measured by UV-Vis spectrophotometer. The correction factor (CF) was determined by measuring A280 and A360 of the pure 100 µM DNP-PEG4-NHS ester solution.

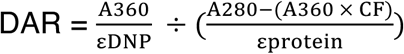

