## Supplemental File for "Low-density lipoprotein receptor-targeting chimeras for membrane protein degradation and enhanced drug delivery"

**This PDF file includes:**

**Extended Data Figures 1-9**

**Extended Table 1**

**References 1-5**

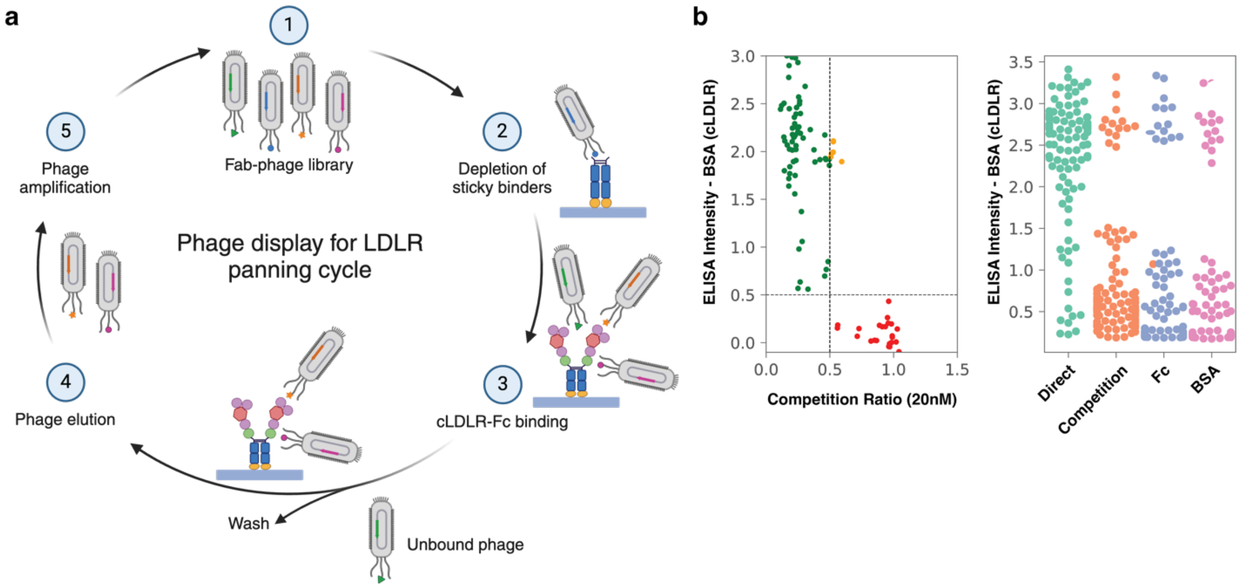

**Extended Data Fig.1. Phage display screening for LDLR-specific antibodies. Related to Figure 1. a**, Phage display workflow for negative and positive selection for cleaved LDLR (cLDLR)-binding Fabs. A Fab-phage library (1) was depleted for binding Fc (2) and residual phage captured on beads that bound the cLDLR-Fc (3) that were washed and eluted (4) and amplified for another round (5). Figure was generated by Biorender. **b**, Phage clones that selectively bound to cLDLR while exhibiting no binding to the negative antigens Fc and bovine serum albumin (BSA), as well as soluble cLDLR in competition assays, were advanced from the ELISA screening for further downstream analysis. Phage clones that passed the screening were highlighted in green dots (upper left quadrant) on left-side diagram.

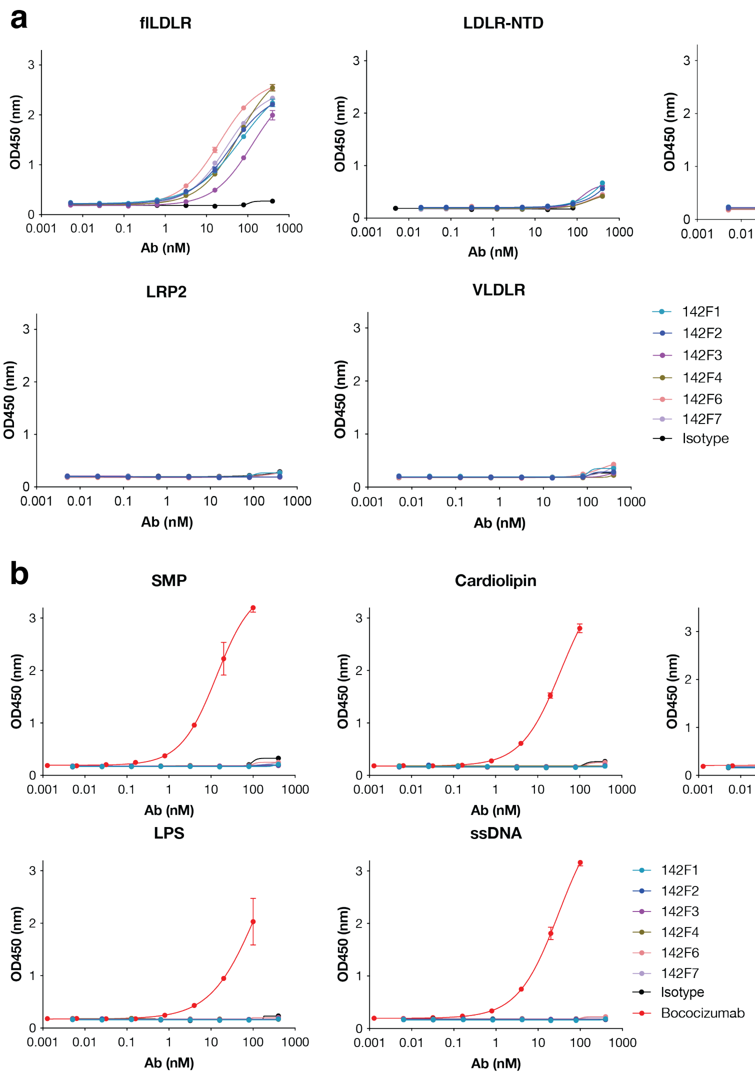

**Extended Data Fig.2. Cross-reactivity and polyspecificity of cLDLR Fabs. Related to Figure 1. a**, ELISA binding of cLDLR Fabs against full-length LDLR (flLDLR), N-terminal ligand-binding domain (NTD) of LDLR, and full-length extracellular domains from LRP8, LRP2, and VLDLR. An anti-CDCP1 Fab 4A06[1], served as a negative control. **b,** ELISA binding of cLDLR Fabs against a panel of commonly used polyspecific reagents[2] including solubilized membrane proteins (SMP), cardiolipin, insulin, LPS, and single-stranded DNA (ssDNA). A polyreactive antibody bococizumab[3] served as a positive control. Each sample was tested in duplicate and error bars represented standard deviations. Data was representative for at least two biological replicates.

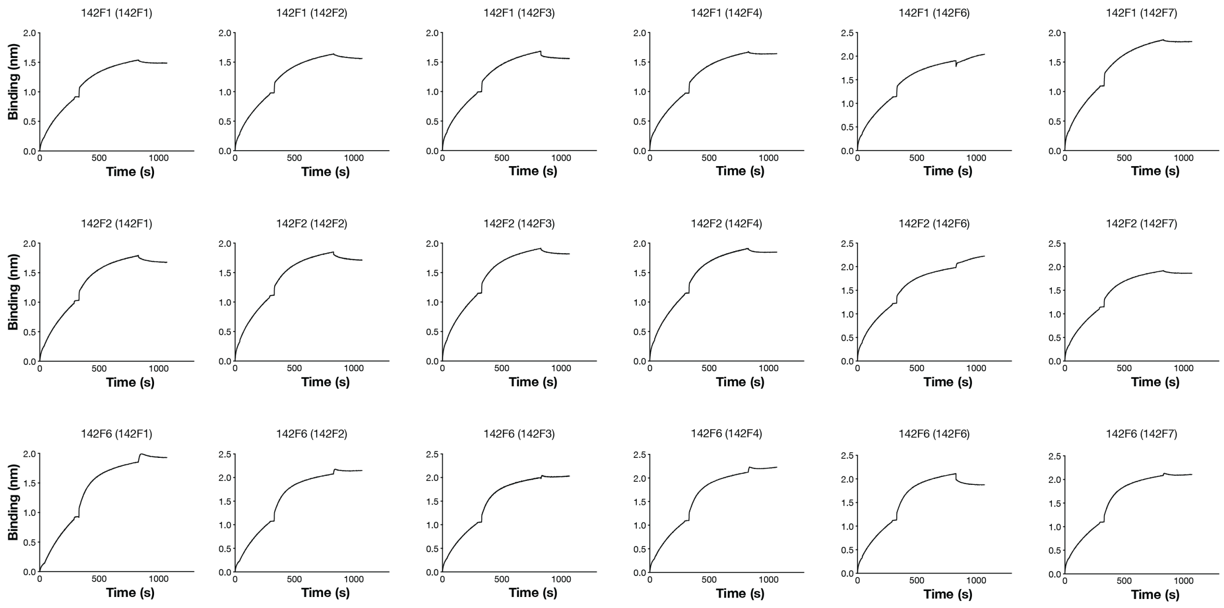

**Extended Data Fig.3. Epitope binning of cLDLR Fabs. Related to Figure 1.** 20 nM of biotinylated cLDLR antigens were captured using streptavidin biosensors. After antigen loading, a saturating concentration of Fab was added for 10 min. Competing concentrations of Fabs (50 nM) were then added for 5 min to measure binding in the presence of saturating antibodies. All incubation steps were performed in 1x PBS + 0.05% Tween + 0.2% BSA at room temperature.

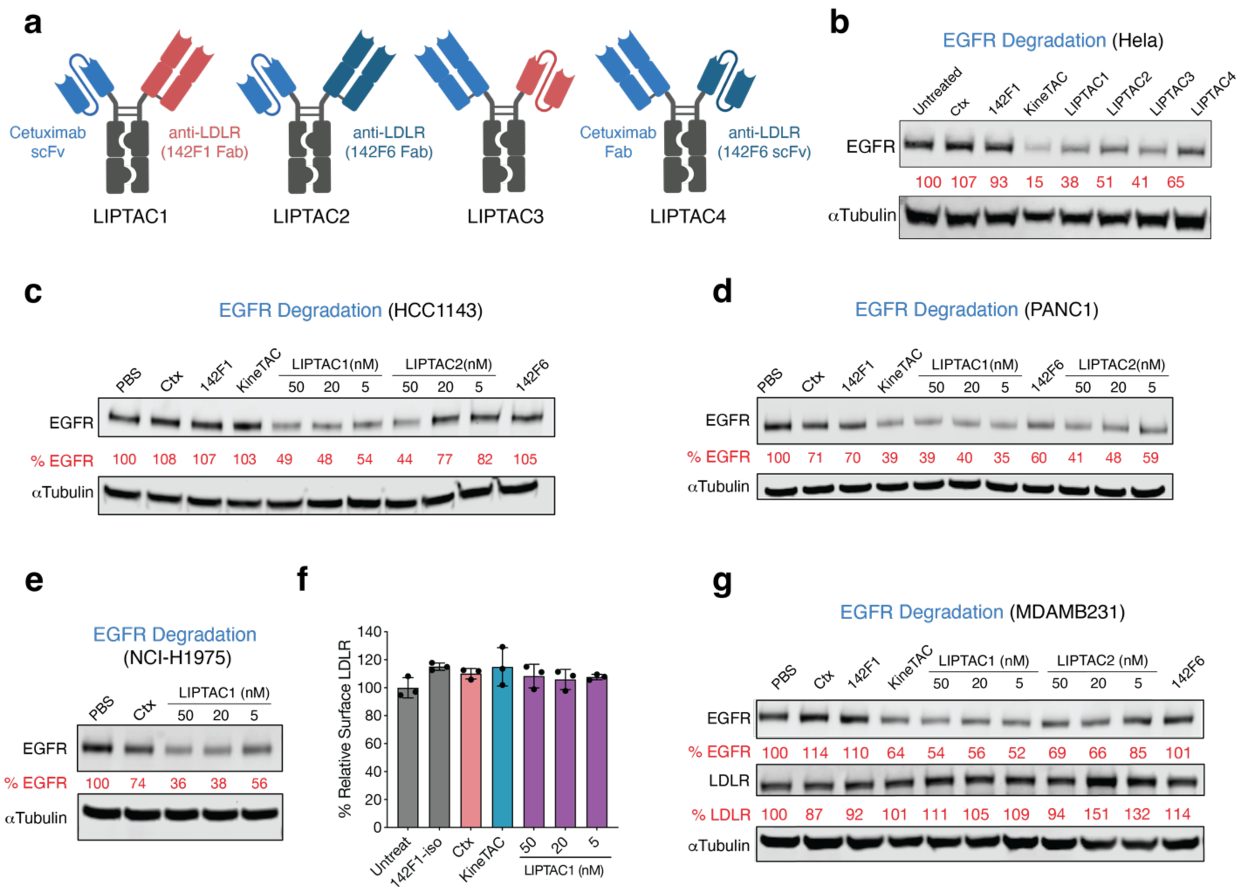

**Extended Data Fig.4. LIPTAC-mediated EGFR degradation in different formats and on multiple cell lines. Related to Figure 2. a**, Schematic of the LDLR-Cetuximab (Ctx) LIPTAC constructs that were tested. **b**, EGFR degradation in HeLa cells after 24 h of treatment with 50 nM Ctx IgG, 142F1 isotype, Ctx-KineTAC, or Ctx-LIPTACs. Data were represented for three independent experiments. **c-e**, Dose escalation experiment showing total EGFR degradation by western blot on HCC1143, PANC-1, and NCI-H1975 cells after 24 h of treatment with Ctx-LIPTACs, or 50 nM of control antibodies. Percent EGFR levels were quantified by ImageJ relative to PBS control. **f**, Flow cytometry analysis showing surface LDLR remained unchanged on MDA-MB-231 cells following 24 h treatment of Ctx-LIPTAC1, 50 nM Ctx-KineTAC, Ctx IgG, or anti-LDLR 142F1 isotype. Percent LDLR was determined by MFI of the APC fluorescence channel of live cells. Each sample was tested in biological triplicate and error bars represented standard deviations. **g**, Western blotting images of EGFR and LDLR on MDA-MB-231 cells following 24 h of antibody treatment. Data were represented for three independent experiments. Percent EGFR levels were quantified by ImageJ relative to PBS control.

| Accession | Statistical significane: log10(p value) | Protein Fold change (log2 LIPTAC/PBS) |
|---|---|---|
| Downregulated | | |
| A1L157\|TSN11_HUMAN | 1.992 | -2.1202942 |
| P55011\|S12A2_HUMAN | 1.649 | -2.0588937 |
| P00533\|EGFR_HUMAN | 9.321 | -1.5563933 |
| Q8NG11\|TSN14_HUMAN | 2.115 | -1.1844246 |
| Q9H2H9\|S38A1_HUMAN | 3.483 | -1.0892673 |
| P43007\|SATT_HUMAN | 6.276 | -1.0291463 |
| Upregulated | | |
| P01130\|LDLR_HUMAN | 8.692 | 1.0071955 |
| Q9UHX3\|AGRE2_HUMAN | 1.563 | 1.02856915 |
| Q9BZ76\|CNTP3_HUMAN | 1.331 | 4.21179098 |
| Q9P1W3\|CSC1_HUMAN | 6.302 | 6 |

**Extended Data Fig.5. Surface proteins on MDA-MB-231 cells having differential abundance with or without LIPTAC1 treatment. Related to Figure 2.** SILAC heavy-labeled cells were treated with PBS control and light-labeled cells were treated with 50 nM bispecific Ctx-LIPTAC1. 48 h after antibody treatment, cells were harvested, cell surface captured, digested, and analyzed by quantitative proteomics. Data are mean of three biological replicates. Surface proteins were annotated by the SURFY database[4]. P values were determined by unpaired two-tailed t tests with PBS-treated cells.

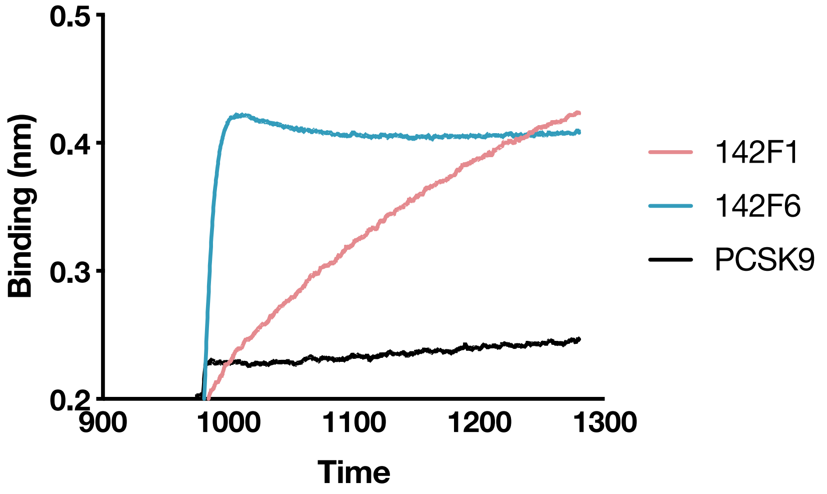

**Extended Data Fig.6. Epitope binning of PCSK9 and anti-LDLR antibodies. Related to Figure 2.** 20 nM of biotinylated cLDLR antigens were captured using streptavidin biosensors. After antigen loading, 200 nM of PCSK9 D374Y protein for 10 min at endosomal pH to improve its binding affinity against LDLR[5]. Competing concentrations of 142F1, 142F6, and PCSK9 (50 nM) were then added for 5 min to measure binding in the presence of saturating antibodies. The curves presented the binding for competing proteins. All incubation steps were performed in 1x PBS + 0.05% Tween + 0.2% BSA at pH 5 at room temperature.

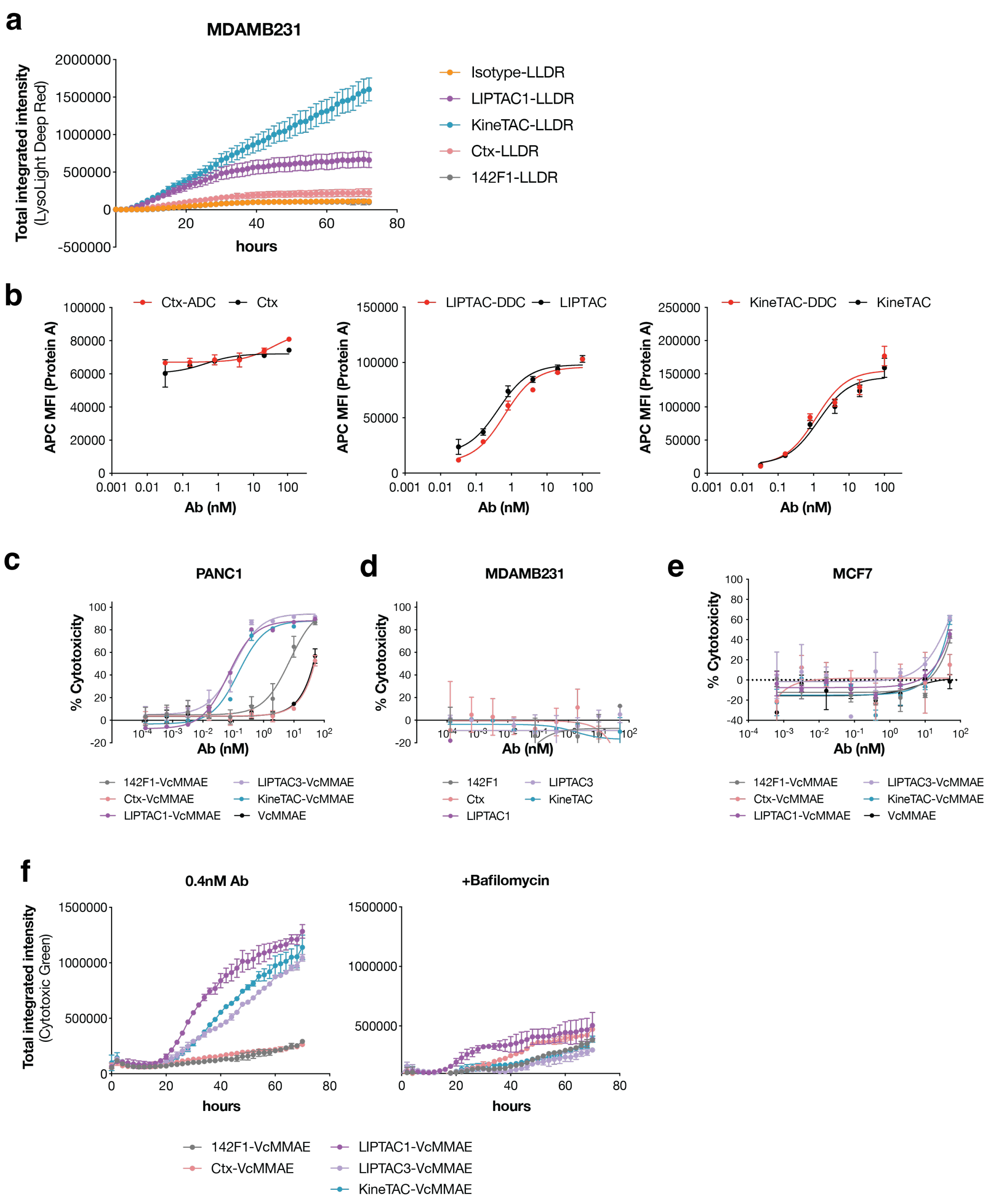

**Extended Data Fig.7. Payload delivery and cytotoxicity of EGFR-targeting DDCs in comparison with ADCs. Related to Figure 4. a**, A payload cleavage assay in EGFR-expressing A431 cells. Cells were treated with 25 nM of antibodies conjugated with lysolight deep red (LLDR) dyes. Images were captured every 2 h for 72 h on the Incucyte. Total integrated intensity was calculated by NIRCU x μm²/image on the Incucyte. Error bars represented standard deviations of at least two biological replicates. **b**, Flow cytometry analysis showing antibodies with or without drug conjugations remained binding to MDA-MB-231 cells. Antibody binding was detected by mean fluorescence intensity (MFI) of AF647-conjugated protein A. Each sample was tested in biological duplicate and error bars represented standard deviations. **c**, Cytotoxicity of Ctx-ADC and Ctx-DDCs on PANC-1 cells. Cell viability was measured after 72 h of incubation using the CellTiter-Glo Reagent. **d**, Cytotoxicity of drug-free antibodies on MDA-MB-231 cells after 72 h of incubation. **e**, Cytotoxicity of Ctx-ADC and Ctx-DDCs after 72 h of incubation on MCF7 that did not express EGFR. Each sample was tested in biological duplicate and error bars represented standard deviations. **f**, Cytotoxicity of 0.4 nM control ADCs and Ctx-DDCs either in LIPTAC or KineTAC formats on A431 cells, with or without the lysosomal inhibitor Bafilomycin A (BafA, 50 nM). Dead cells were labeled by a cytotoxic green dye. Total integrated intensity was calculated by GreenCU x µm2/image on the Incucyte and the images were captured every 2 h for a total of 72 h.

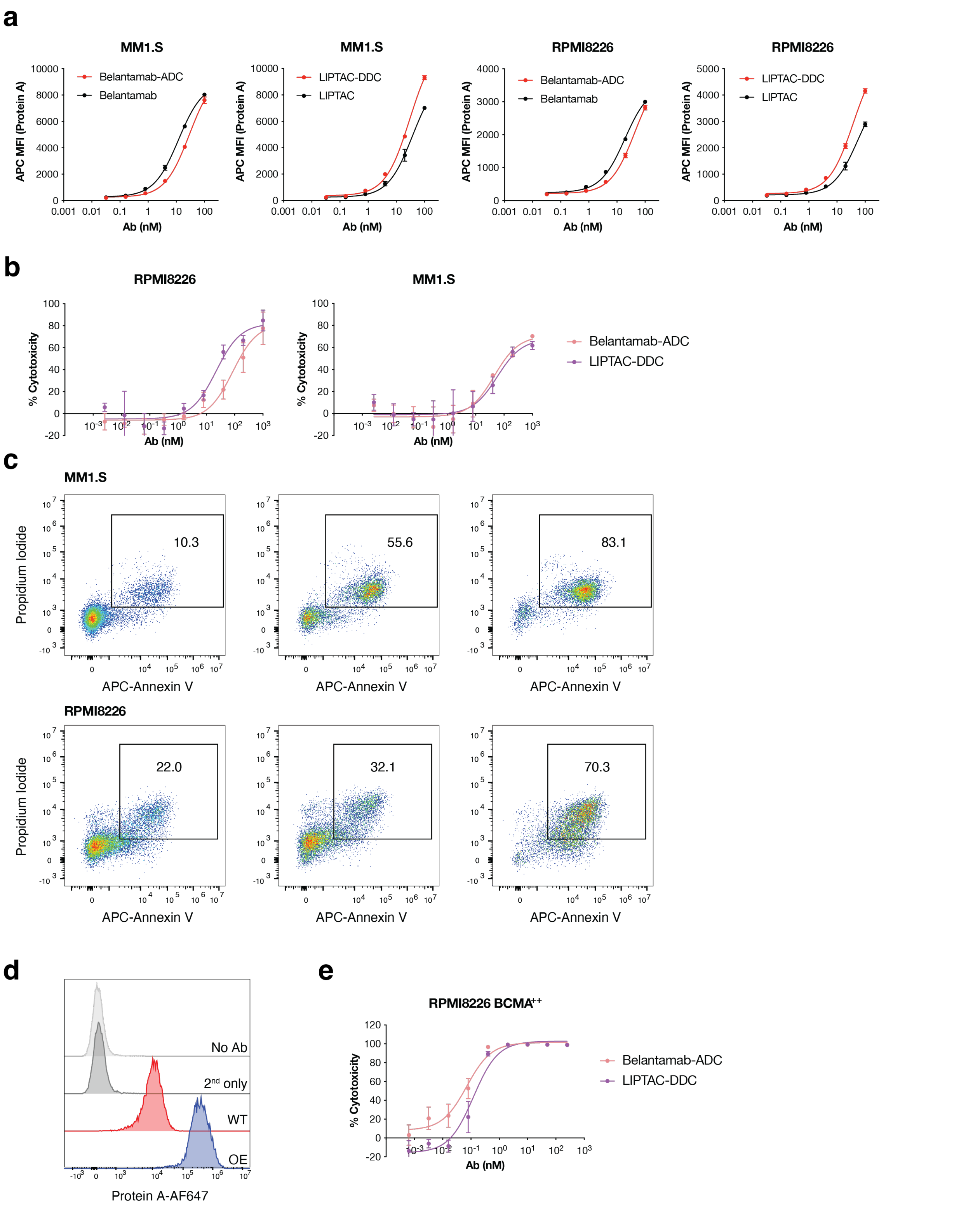

**Extended Data Fig.8. Cytotoxicity of BCMA-targeting ADC and DDC. Related to Figure 5. a**, Flow cytometry analysis of drug-conjugated antibody and drug-free antibody binding to MM1.S and RPMI8226 cells. Cells were gated from live singlets. Antibody binding was detected by mean MFI of AF647-conjugated protein A. Each sample was tested in biological duplicate and error bars represented standard deviations. **b**, Cytotoxicity of Belantamab-ADC and LIPTAC-DDC on RPMI8226 as well as MM1.S cells. Cell viability was measured after 96 h of incubation using the CellTiter-Glo Reagent. Each sample was tested in biological triplicate and error bars represent standard deviations. **c**, Flow cytometry-based cell viability assay in MM1.S and RPMI8226 cells after 4 days treatment with 31.25 nM Belantamab-ADC or LIPTAC-DDC. Cells were stained with propidium iodide and APC-annexin V for 30 min on ice in the presence of 1 mM calcium chloride, then washed twice with PBS + 1% BSA. **d**, Flow cytometry histogram analysis of wildtype (WT) and BCMA overexpressed (OE) RPMI8226 cells. Cells were stained with Belantamab first, followed by protein A-AF647 secondary staining. **e**, Cytotoxicity of Belantamab-ADC and LIPTAC-DDC on RPMI8226 as well as BCMA overexpressing RPMI8226 cells. Each sample was tested in technical triplicate and error bars represented standard deviations. Data is representative for two biological replicates.

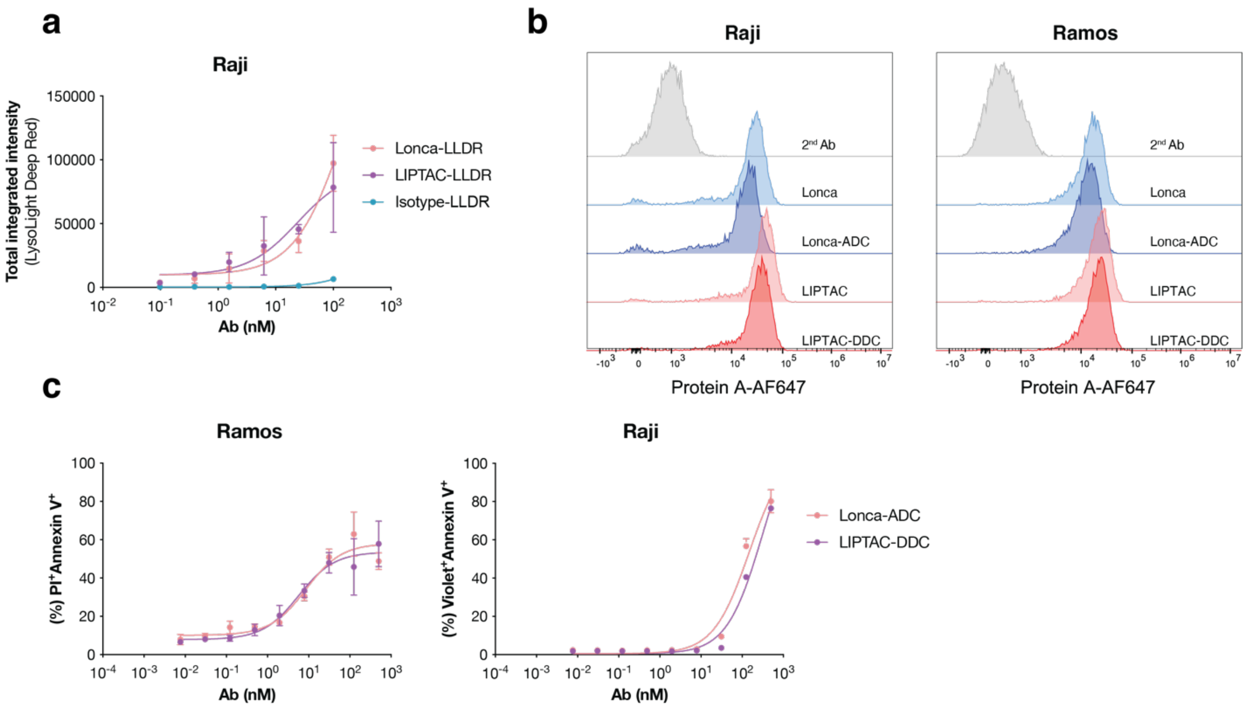

**Extended Data Fig.9. Cytotoxicity of CD19-targeting ADC and DDC. Related to Figure 5. a**, Antibody internalization assay in Raji cells after 72 h treatment with LLDR-labeled antibodies. Each sample was tested in duplicate and error bars represented standard deviations. Total integrated intensity was calculated by NIRCU x µm$^2$/image on the Incucyte. **b**, Flow cytometry analysis of drug-conjugated antibody and drug-free antibody binding to Raji and Ramos cells. Antibody binding was detected by an AF647-conjugated protein A. **c**, Flow cytometry-based cell viability assay in Ramos and Raji cells after 4 days treatment with Loncastuximab (Lonca)-ADC or LIPTAC-DDC. Cells were stained with propidium iodide (PI) and APC-annexin V for 30 min on ice in 1 mM calcium chloride, then washed twice with PBS + 1% BSA. Each sample was tested in biological duplicate and error bars represented standard deviations.

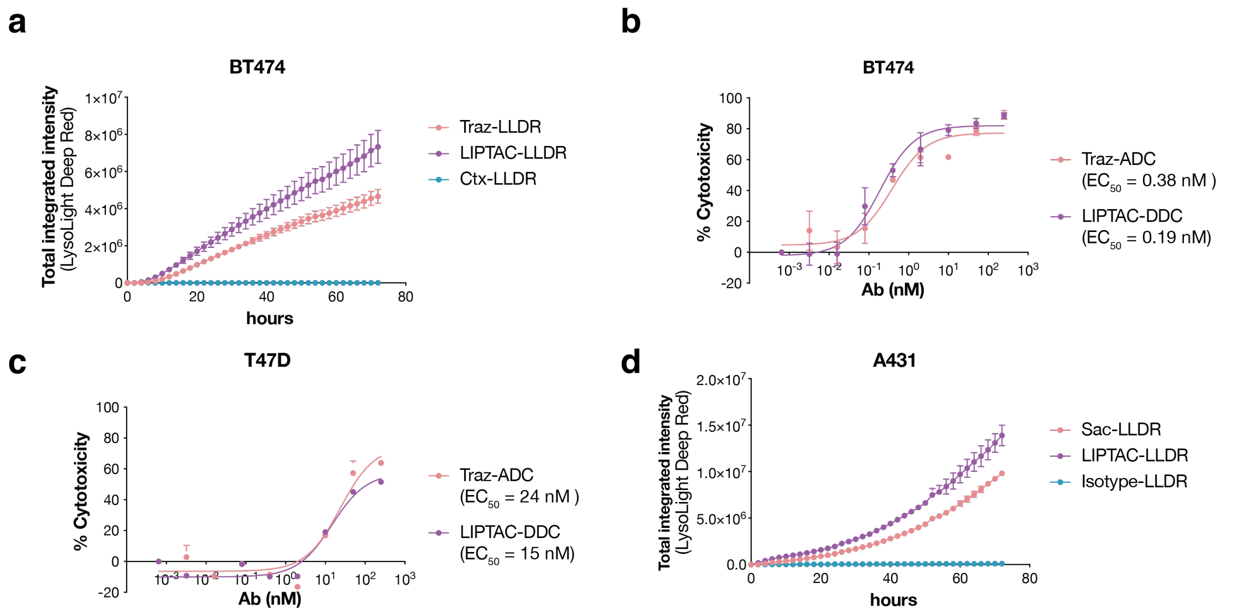

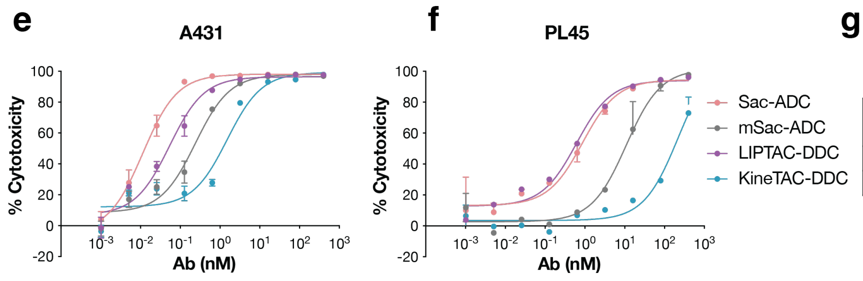

**g**

| $EC_{50}$ (nM) | Sac-ADC | mSac-ADC | LIPTAC DDC | KineTAC -DDC |
|---|---|---|---|---|
| A431 | 0.01 | 0.24 | 0.05 | 1.5 |
| PL45 | 0.87 | 11 | 0.62 | 220 |

**Extended Data Fig.10. Cytotoxicity of HER2-targeting and TROP2-targeting ADCs and DDCs. Related to Figure 5. a**, Antibody internalization assay in BT474 cells treated with 100 nM of LLDR-labeled antibodies. Each sample was tested in triplicate and error bars represented standard deviations. Images were captured every 2 h for 72 h on the Incucyte. Total integrated intensity was calculated by NIRCU x µm$^2$/image on the Incucyte. **b**,**c**, Cytotoxicity of Trastuzumab (Traz)-ADC and LIPTAC-DDC on HER2$^{high}$ BT474 and HER2$^{low}$ T47D cells. Cell viability was measured after 72 h of incubation using the CellTiter-Glo Reagent. Each sample was tested in biological duplicate and error bars represented standard deviations. **d**, Antibody internalization assay in A431 cells treated with 100 nM of LLDR-labeled antibodies. Each sample was tested in triplicate and error bars represented standard deviations. Images were captured every 2 h for 72 h on the Incucyte. Total integrated intensity was calculated by NIRCU x µm$^2$/image on the Incucyte. **e**,**f**, Cytotoxicity of Sacituzumab (Sac)-ADC, monomeric Sacituzumab (mSac)-ADC, LIPTAC-DDC and KineTAC-DDC on TROP2$^{high}$ A431 and TROP2$^{medium}$ PL45 cells. Cell viability was measured after 72 h of incubation using the CellTiter-Glo Reagent. Each sample was tested in triplicate and error bars represented standard deviations. **g**, $EC_{50}$ table for cytotoxicity of TROP2-targeting ADCs and DDCs summarized from **e,f**. $EC_{50}$ values were calculated using "One-Site Fit LogIC50" regression in GraphPad Prism 10.2.

| Labeled Antibody | A280 | A360 | DAR |
| --- | --- | --- | --- |
| Ctx-DNP | 5.72 | 0.73 | 1.85 |
| LIPTAC1-DNP | 1.95 | 0.45 | 2.70 |
| LIPTAC3-DNP | 0.52 | 0.1 | 2.22 |
| KineTAC-DNP | 2.32 | 0.66 | 2.85 |
| 142F1-Iso-DNP | 6.4 | 2.09 | 3.45 |

Average DAR = 2.62

Correction factor = A280/A360 = 0.445/1.41 = 0.315

**Extended Table 1. Characterization of relative DAR. Related to Figure 4.** Antibodies were labeled with DNP-PEG4-NHS ester for 2 h at RT using the same method as NHS ester-PEG4-ValCit-PAB-MMAE labeling. The protein and DNP absorbances were measured at 280 nm and 360 nm respectively. Correction factor was measured by the absorbance of pure DNP-PEG4-NHS ester at 280 nm and 360 nm respectively. Each sample was tested in technical duplicate.
